# *Kluyveromyces marxianus* as a robust synthetic biology platform host

**DOI:** 10.1101/353680

**Authors:** Paul Cernak, Raissa Estrela, Snigdha Poddar, Jeffrey M. Skerker, Ya-Fang Cheng, Annika K. Carlson, Berling Chen, Victoria M. Glynn, Monique Furlan, Owen W. Ryan, Marie K. Donnelly, Adam P. Arkin, John W. Taylor, Jamie H. D. Cate

## Abstract

Throughout history, the yeast *Saccharomyces cerevisiae* has played a central role in human society due to its use in food production and more recently as a major industrial and model microorganism, because of the many genetic and genomic tools available to probe its biology. However *S. cerevisiae* has proven difficult to engineer to expand the carbon sources it can utilize, the products it can make, and the harsh conditions it can tolerate in industrial applications. Other yeasts that could solve many of these problems remain difficult to manipulate genetically. Here, we engineer the thermotolerant yeast *Kluyveromyces marxianus* to create a new synthetic biology platform. Using CRISPR-Cas9 mediated genome editing, we show that wild isolates of *K. marxianus* can be made heterothallic for sexual crossing. By breeding two of these mating-type engineered *K. marxianus* strains, we combined three complex traits– thermotolerance, lipid production, and facile transformation with exogenous DNA-into a single host. The ability to cross *K. marxianus* strains with relative ease, together with CRISPR-Cas9 genome editing, should enable engineering of *K. marxianus* isolates with promising lipid production at temperatures far exceeding those of other fungi under development for industrial applications. These results establish *K. marxianus* as a synthetic biology platform comparable to *S. cerevisiae*, with naturally more robust traits that hold potential for the industrial production of renewable chemicals.

## SIGNIFICANCE

The yeast *Kluyveromyces marxianus* grows at high temperatures and on a wide range of carbon sources, making it a promising host for industrial biotechnology to produce renewable chemicals from plant biomass feedstocks. However, major genetic engineering limitations have kept this yeast from replacing the commonly-used yeast *Saccharomyces cerevisiae* in industrial applications. Here, we describe genetic tools for genome editing and breeding *K. marxianus* strains, which we use to create a new thermotolerant strain with promising fatty acid production. These results open the door to using *K. marxianus* as a versatile synthetic biology platform organism for industrial applications.

## INTRODUCTION

Synthetic biology is used to harness the metabolic capacity of microorganisms for the biosynthesis of simple and complex compounds now sourced unsustainably from fossil fuels, or that are too expensive to make using chemical synthesis at industrial scale. The yeast *Saccharomyces cerevisiae* has served as the major eukaryotic organism for synthetic biology, but lacks the metabolic potential that could be exploited in many of the more than one thousand yeast species that have been identified to date. These yeasts remain difficult to use, however, as there are few synthetic biology tools to access their underlying metabolic networks and physiology (1). The budding yeast *Kluyveromyces marxianus* possesses a number of beneficial traits that make it a promising alternative to *S. cerevisiae. K. marxianus* is the fastest growing eukaryotic organism known (2), is thermotolerant, growing and fermenting at temperatures up to 52 °C and 45 °C, respectively (3) and uses a broad range of carbon sources, including pentose sugars. These traits are polygenic and would be difficult to engineer into a less robust host such as *S. cerevisiae. K. marxianus* also harbors high strain-to-strain physiological and metabolic diversity, which could prove advantageous for combining beneficial traits by sexual crossing. However, *K. marxianus* is generally found to be homothallic (4, 5) (i.e. is self fertile), and cannot be crossed in a controlled manner.

For *K. marxianus* to be useful as a yeast platform for synthetic biology, it will be essential to establish efficient gene editing tools along with methods to cross strains with stable ploidy. These tools would enable rapid strain development, by generating genetic diversity and facilitating stacking of industrially important traits. The CRISPR-Cas9 gene editing system has been used in many yeasts including *S. cerevisiae, S. pombe, Y. lipolytica, K. lactis* and recently *K. marxianus* (6–9). Genome editing in *K. marxianus* should allow manipulation of known genetic targets. However, most desired traits likely depend on multiple, unlinked genetic loci, which remain difficult to identify without the ability to carry out genetic crosses. The ability to cross phenotypically diverse *S. cerevisiae* strains has been an indispensable tool for exploring its biology on a genome-wide scale, and for improving its use as an industrial host (10). To approach the versatility of *S. cerevisiae* genetics, it will be necessary to gain full control over *K. marxianus* ploidy and mating type. As a homothallic yeast, *K. marxianus* lacks a permanent mating type, because *K. marxianus* haploid cells naturally change their mating type (either the **a**-mating type, *MAT***a**, or α-mating type, *MAT*α) leading to uncontrolled *MAT***a***/MAT*α diploidization within a population (11). This stochastic ploidy makes it impossible to carry out quantitative biological studies of interesting traits that are ploidy-specific (12), as it leads to populations with mixed phenotypes, and prevents *K. marxianus* domestication through selective crossing.

To overcome the limitations in using *K. marxianus* as a synthetic biology platform, we adapted the CRISPR-Cas9 system we developed for *S. cerevisiae* (13) for use in *K. marxianus*, enabling both non-homologous end joining (NHEJ) and homology directed repair (HDR) based genome editing. We identified the genetic loci responsible for mating type switching in *K. marxianus* and created domesticated laboratory strains by simultaneously inactivating these genes. With this platform in place, we explored a large collection of wild *K. marxianus* strains to investigate *K. marxianus* lipid production at high temperatures. By domesticating and crossing promising strains, we were able to combine three complex traits-the ability to uptake exogenous DNA (transformability), thermotolerance and higher lipid production-into single *K. marxianus* isolates.

## RESULTS

### CRISPR-Cas9 system in *K. marxianus*

We established robust genome editing in *K. marxianus* by adapting the plasmid-based CRISPR-Cas9 (CRISPRm) system we previously developed for *S. cerevisiae* (13). We first identified a *K. marxianus-specific* origin of replication and *K. marxianus-*specific promoters and terminators for expressing Cas9 (*SI Materials and Methods*). We used the *S. cerevisiae* gene for tRNA^Phe^ as an RNA polymerase III promoter to express single-guide RNAs (sgRNAs) from the same plasmid (**Fig. 1*A***). To test the effectiveness of the redesigned CRISPR-Cas9 system, we used a wild strain we isolated from a sugarcane bagasse pile (*Km*1, **Table 1**), and a *Km*1-derived *MAT***a** heterothallic strain (*Km*30, *SI Appendix*, Table S1). We transformed the pKCas plasmid (G418^R^) encoding an sgRNA targeting the *URA3* gene into *Km*1, and selected for 5-fluoroorotic acid (5-FOA) resistance to identify *ura3*^−^ colonies. The efficiency of NHEJ-based Cas9 editing (CRISPR_NHEJ_)-the number of *ura3*^−^ colonies divided by the number of total G418^R^ transformants-was near 90%, and was confirmed by sequencing the *URA3* locus. Targeting other genes across the genome resulted in around 75% efficiency (*SI Appendix*, Fig. S1). To test the ability of the *K. marxianus* CRISPR system to insert exogenous DNA at a defined locus, we also co-transformed into strain *Km*30 the pKCas plasmid encoding a guide RNA targeting the *URA3* gene along with a double-stranded DNA repair template comprised of a linear Nourseothricin resistance cassette flanked by *K. marxianus URA3* homology sequences adjacent to the Cas9 target site (NatMX flanked by 0.9 Kb homology arms). Using replica plating of G418^R^ transformants onto two selection plates, 5-FOA to detect *ura3*^−^ alleles, and Nat^R^ to detect HDR events, we found 100% of the colonies to be *ura3*^−^, and ~97% (189/195) to be Nat^R^, indicating that repair of Cas9-induced double strand breaks allowed highly-efficient HDR-mediated gene integration (CRISPR_HDR_). We used colony PCR on select Nat^R^ colonies (targeting outside the 0.9 kb homology arms of the NatMX cassette), to confirm NatMX cassette integration at the *URA3* locus.

**Table 1.** List of wild type strains used in this work.

| **Strain** | **ATCC** | **NCYC** | **CBS** | **NRRL** |
| --- | --- | --- | --- | --- |
| *Km1\** |  |  |  |  |
| *Km2* | 10022 | 100 | 6432 | Y-665 |
| *Km5* | 46537 | 851 | 397 | Y-2415 |
| *Km6* |  | 143 | 608 | Y-8281 |
| *Km9* | 26548 | 2597 | 6556 | Y-7571 |
| *Km11* | 36907 | 587 |  |  |
| *Km16* | 10606 |  | 396 | Y-1550 |
| *Km17* | 8635/28910 |  |  | Y-1190 |
| *Km18* |  |  | 1089 | Y-2265 |
| *Km19* | 26348 |  |  |  |
| *Km20\** |  |  |  |  |
| *Km21\** |  |  |  |  |

\**Km1*, *Km20* and *Km21* were isolated from sugarcane bagasse piles. *Km1* was isolated at Raceland Raw Sugar Corporation, Raceland, Louisiana. *Km20* and *Km21* were isolated at the Sugarcane Growers' Cooperative, Belle Glade, Florida.

**Figure 1.**
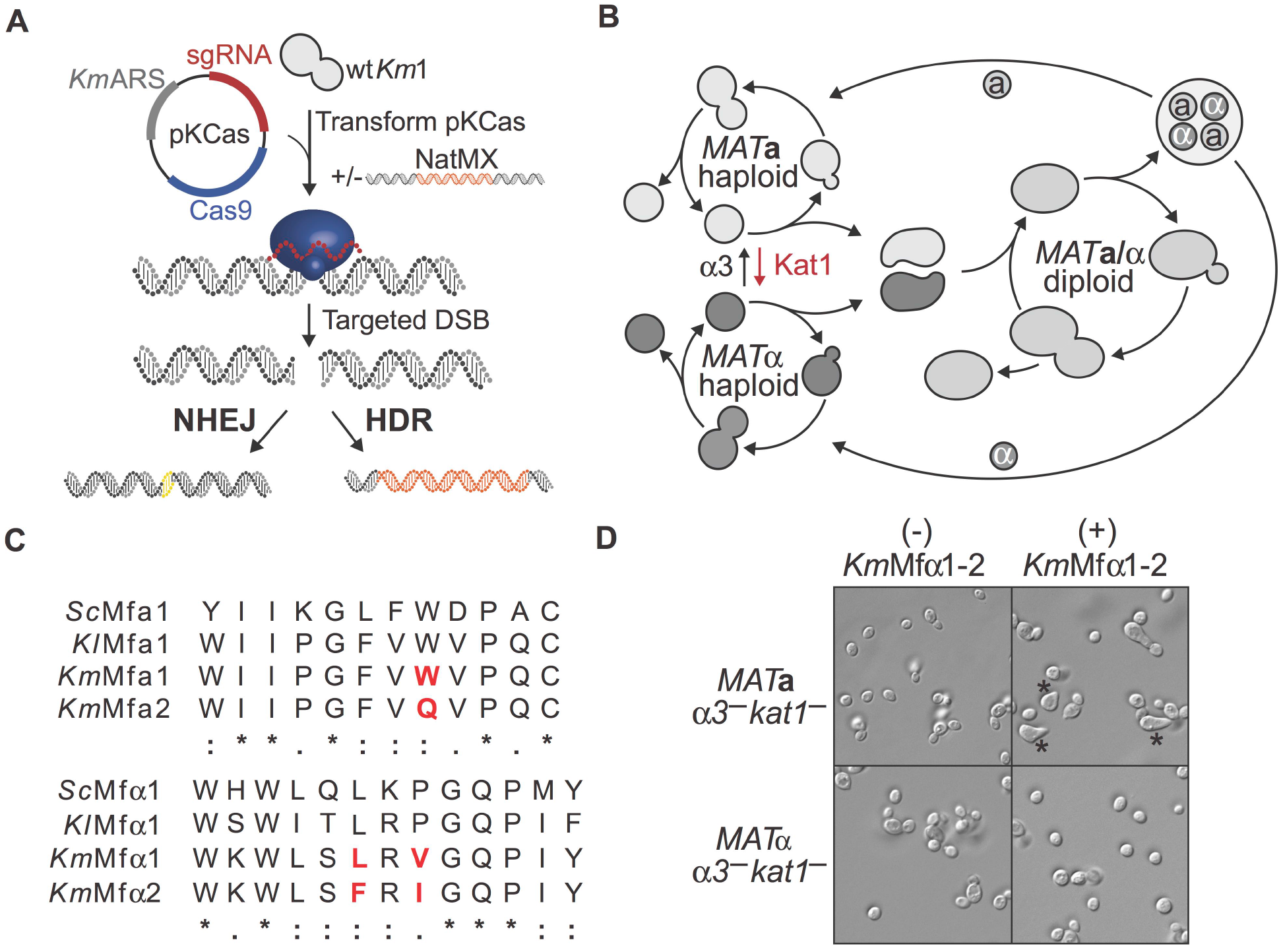
Genetic manipulation of *K. marxianus* strains. (**A**) CRISPR_NHEJ_ and CRISPR_HDR_ systems. *K. marxianus* transformed with the pKCas plasmid generates small indels near the cut site, a common product of non-homologous end joining (NHEJ) repair of the DNA double strand break. When transformed with both the pKCas plasmid and a donor DNA, homologous recombination products are seen in the target site. (**B**) Yeast life cycle. Haploid *MAT***a** and *MAT*α switch mating type by transposases α3 and Kat1 in *K. lactis.* Haploid cells conjugate to form *MAT***a**/*MAT*α diploids. Diploids undergo meiosis to form haploid spores that germinate to complete the life cycle. (**C**) Mature **a**- and α-pheromones from *K. marxianus* aligned with the *S. cerevisiae* and *K. lactis* sequences. Red, non-conserved amino acids between *K. marxianus* **a**- and α-factors. Amino acids are marked as identical (*), with similar polarity (:), or with different polarity (.). (**D**) Putative heterothallic *MAT***a** and *MAT*α strains incubated with a cocktail of both mature α-factor pheromones (*Km*Mf α1-2) results in mating projections from the MAT**a** strain only (*).

### Engineering mating-competent heterothallic *K. marxianus* strains

We used CRISPR_NHEJ_ to make stable *K. marxianus* laboratory strains with defined ploidy and mating type to enable the use of classical yeast genetics. Most naturally-isolated *K. marxianus* strains are homothallic, i.e. they change their mating type spontaneously by “mating-type switching” to create mixed populations of *MAT***a**, *MAT*α and *MAT***a**/*MAT*α cells (4, 5). The *K. marxianus* mating type switching mechanism is not genetically conserved with the well-characterized HO endonuclease mechanism employed by *S. cerevisiae.* Notably, a two-component switching mechanism has been identified in *Kluyveromyces lactis* (14, 15), which uses two transposases (Kat1 and α3) for *MAT* switching. The α3 transposase switches *MAT*α type cells to *MAT***a** type and Kat1 switches the *MAT***a** to *MAT*α type (**Fig. 1*B***).

We identified the *K. marxianus* orthologs of the *K. lactis KAT1* and *ALPHA3* genes using reciprocal BLASTp against predicted ORFs from the whole genome sequence of *Km*1 (*SI Appendix*, Table S2) (16). Using CRISPR_NHEJ_, we targeted both transposase genes (*SI Appendix*, Table S3) to create frameshift mutation loss-of-function alleles. We then isolated several of these double-transposase inactivated *Km*1 α3^−^ *kat1*^−^ strains that had small base pair insertions or deletions near the Cas9 cut site (*SI Appendix*, Fig. S1). To identify *MAT***a** haploid isolates, we used a pheromone morphological response assay. Yeast mating is initiated by the secretion of small peptide pheromones **a**-factor and α-factor by *MAT***a** and *MATα* cells, respectively. The pheromones, derived from **α**-pheromone and α-pheromone precursor proteins, **m**ating **f**actor **a** (MFA1 and MFA2) and mating factor α (MFα1), are detected by their cognate cell surface recognition proteins and lead to polar morphogenesis or the formation of mating projections (“shmoo”) that can be used to deduce a strain’s mating type. We identified two putative *MFA* genes (*KmMFA1* and *KmMFA2*) as well as the *MF*α gene (*Km*MFα1, encoding two isotypes, *Km*MFα1 and *Km*MFα2) in the *K. marxianus* genome by reciprocal BLASTp using the *S. cerevisiae* and *K. lactis* protein sequences as queries (16, 17) (**Fig. 1*C***) (*SI Appendix*, Fig. S2). Incubation of *K. marxianus* strain *Km*1 α*3*^−^ *kat1*^−^ cells with synthetic *Km*MFα1 and *Km*MFα2 peptide pheromones resulted in isolates that responded to both α-factors (**Fig. 1*D***), indicating these are *MAT***a** α*3*^−^ *kat1*^−^ haploids. We categorized unresponsive strains as either *MAT*α or diploid strains, using sequencing of the *MAT* locus (Fig. S3).

Once we identified stable *MAT***a** α*3*^−^ *kat1*^−^ or *MAT*α α*3*^−^ *kat1*^−^ strains through the pheromone response method and subsequent sequencing of the *MAT* locus (*SI Appendix*, Table S2 and Fig. S3), we tested for the ability of two haploid strains with different auxotrophic markers to mate and form prototrophic diploids. First, we used CRISPR_NHEJ_ to create *leu2*^−^ or *trp1*^−^ auxotrophic mutants of the predicted heterothallic *MAT***a** α*3*^−^ *kat1*^−^ and *MAT*α α*3*^−^ *kat1*^−^ or *MAT***a**/*MAT*α strains. Then, *MAT***a** α*3*^−^ *kat1*^−^ *trp1*− and *MAT*α α*3*^−^ *kat1*^−^ *leu2*^−^ strains were combined by opposing streaks on mating-inducing medium. After 2 days, successful mating of *MAT***a** α*3*^−^ *kat1*^−^ *trp1*^−^ and *MAT*α α*3*^−^ *kat1*^−^ *leu2*^−^ cells resulted in growth when cells were replicated plated onto minimal medium lacking tryptophan and leucine (**Fig. 2*A***). To test whether the engineered heterothallic strains could still mate with homothallic wild-type isolates, heterothallic *Km*1 *MAT***a** α*3*^−^ *kat1*^−^ *trp1*^−^ cells were streaked with the homothallic haploid *Km*1 *leu2*^−^ strain resulting in mating and diploid growth. These data confirm the CRISPR_NHEJ_ engineered *Km* α*3*^−^ *kat1*− strains are heterothallic and result in stable haploid, breeding-competent isolates that can mate with opposite mating types and wild homothallic strains.

**Figure 2.**
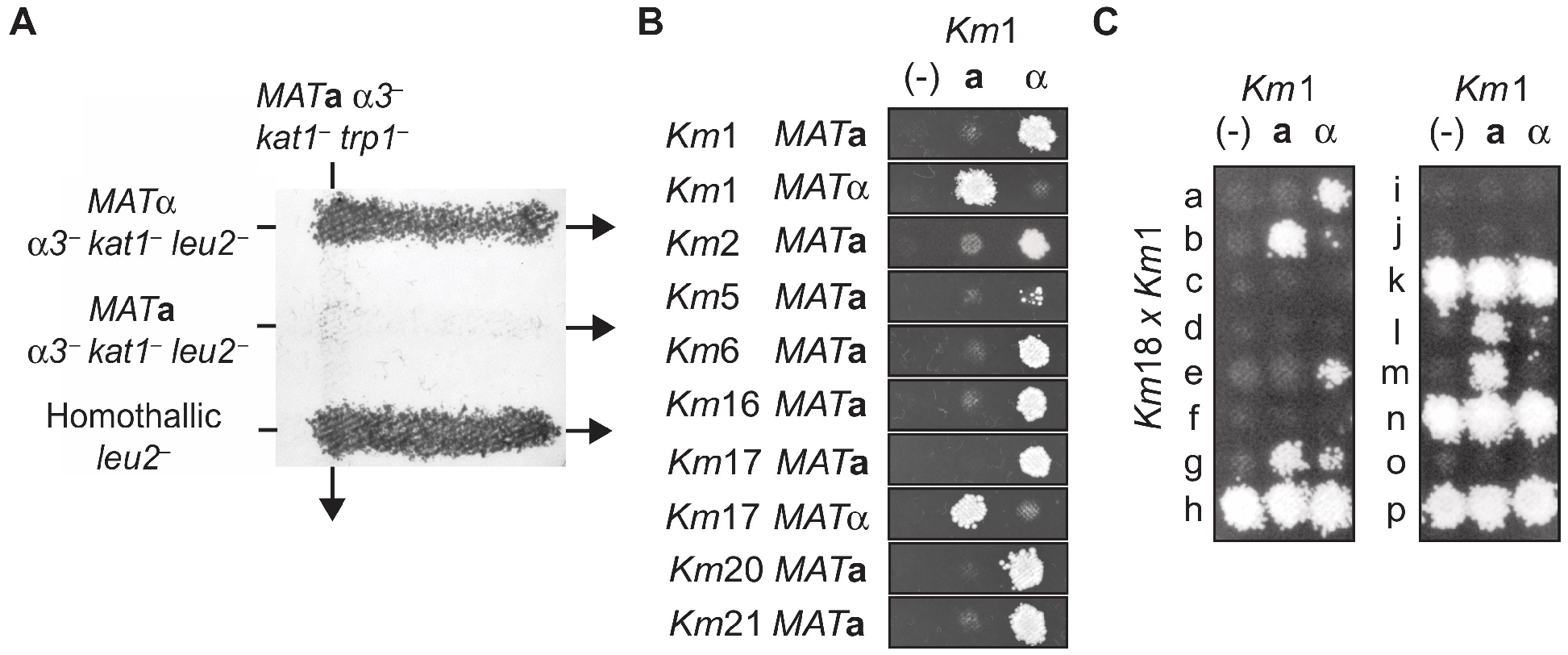
Creation of heterothallic *K. marxianus* strains. **(A)** Auxotrophic mating assay of *Km*1 strains. Strains *Km*1 *MAT*α *α3*^−^ *kat1*^−^ *leu2*^−^, *Km*1 MAT**a** *α3*^−^ *kat1*^−^ *leu2*^−^ and homothallic *Km*1 *leu2*^−^, streaked through strain *Km*1 *MAT***a** *α3*^−^ *kat1*^−^ *trp1*^−^ on 2% glucose plates and replica plated onto SCD minus (Leu, Trp) plates after 2 days. Diploid growth is seen only upon sexual crossing between strains with opposite mating-types or with homothallic haploid strains. **(B)** Auxotrophic mating assay of several triple-*α3*^−^ *kat1*^−^ *leu2*^−^ inactivation strains and *Km*1 *MAT***a** *α3*^−^ *kat1*^−^ *trp1*^−^ or *Km*1 *MAT*α *α3*^−^ *kat1*^−^ *trp1*^−^. Putative heterothallic strains spotted over (-), negative control; **a**, *Km*1 *MAT***a** α*3*^−^ *katT*^−^ *trp1*^−^ reference; **a**, *Km*1 *MAT***a** *α3*^−^ *kat1*^−^ trp1^−^ reference on glucose plates for mating. Replica plating onto SCD minus (Leu, Trp) results in diploid growth. **(C)** The wild homothallic isolate *Km*18 was made *trp*^−^ by UV mutagenesis and crossed with heterothallic *Km*1 *MAT***a** *α3*^−^ *kat1*^−^ *leu2*^−^. Diploids were sporulated, 16 spores isolated (a-p), germinated and the resulting haploids were screened for heterothallic strains by crossing with *Km*1 *MAT***a** *α3*^−^ *kat1*^−^ *trp1*^−^ or *Km*1 *MAT*α *α3*^−^ *kat1*^−^ *trp1*^−^. Screened haploids were auxotrophic strains unable to mate (c, d, f, i, j, o), possible *trp*^−^ revertants (h, k, n, p), homothallic (g) or heterothallic (a, b, e, l, m).

To further validate the role of α3 and Kat1 in switching mating types, we performed complementation assays by constitutively expressing these transposases from plasmids. Plasmids encoding Kat1 or α3 were transformed into *Km*1 α*3*^−^ *kat1*^−^ *leu2*^−^ strains to revert the controlled mating phenotype and promote homothallism. Transformants were then tested for mating-type switching by crossing them with stable heterothallic reference strains (α*3*^−^ *kat1*^−^ *trp1*^−^) of either *MAT***a** or *MAT*α mating type. Using the auxotrophic mating assay, these experiments showed that complementing stable *MAT***a** mutants with Kat1 overexpression plasmids or stable *MAT*α mutants with α3 expressing plasmids induced mating-type switching. Kat1 caused *MAT***a** isolates to switch and mate with a *MAT***a** reference strain, and α3 caused *MAT*α isolates to switch and mate with a *MAT*α reference strain (*SI Appendix*, Fig. S4).

A strong advantage of turning *K. marxianus* into a synthetic biology chassis for metabolic engineering is its high strain-to-strain phenotypic and metabolic diversity (18). To build a widely-useful yeast platform, we sought to create heterothallic strains of each mating type for 12 wild isolates collected from the ATCC, CBS culture collections and our own isolates (**Table 1**). These strains have been isolated from diverse locations and substrates around the world, from dairy to sugarcane bagasse. We used CRISPR_NHEJ_ to create strains with inactivation of three genes, those encoding the transposases Kat1, α3 and an auxotrophic marker, either *TRP1* or *LEU2*. Triple inactivation strains (α*3*^−^ *kat1*^−^ *leu2*^−^ or α*3*^−^ *kat1*^−^ *trp1*^−^) were successfully isolated from 10 of the isolates. We assayed these strains for mating type by crossing them with heterothallic *Km*1 strains as a reference, using the auxotrophic mating assay described above. Heterothallic haploids (*MAT***a** and/or *MAT*α) were isolated from 10 of the triple inactivation strains (**Fig. 2*B***). For strain *Km*18, which was difficult to transform with plasmid DNA, stable heterothallic strains could be isolated from a cross between a homothallic *Km*18 strain first made *trp*^−^ using UV mutagenesis and *Km*1 heterothallic strains. The *Km*18 *trp*^−^ × *Km*1 diploids were sporulated and germinated and then back-crossed with *Km*1 haploid reference strains to establish their mating type (**Fig. 2*C***).

### *K. marxianus* strains engineered for higher levels of lipogenesis

To explore the industrial potential of *K. marxianus* compared to *S. cerevisiae* (i.e. thermotolerance and Crabtree-negative growth, preferring respiration over fermentation (19)), we tested lipid production in *K. marxianus* in aerobic conditions. We first screened 11 wild-type *K. marxianus* isolates for levels of lipogenesis using a lipophilic fluorescent dye (Nile red), combined with flow cytometry and cell sorting. Nile red localizes to lipid droplets in yeast and exhibits increased red fluorescence proportional to the total amount of lipid in the cell (20, 21). The *K. marxianus* strains were grown in 8% glucose or 8% cellobiose lipogenesis medium at 30 °C and 42 °C, and time points were collected every 24 hours to be analyzed by flow cytometry. The highest fluorescence was observed in strains fed 8% glucose at 42 °C for 24 h, with large strain-to-strain differences spanning a ~20-fold change in fluorescence (**Fig. 3*A***). A few strains also produced significant amounts of lipid in cellobiose at 42 °C, when compared to their production of lipid in glucose (i.e. strains *Km*2 and *Km*17, **Table 1**) (*SI Appendix*, Fig. S5). All strains produced much lower levels of lipid when grown at 30 °C.

**Figure 3.**
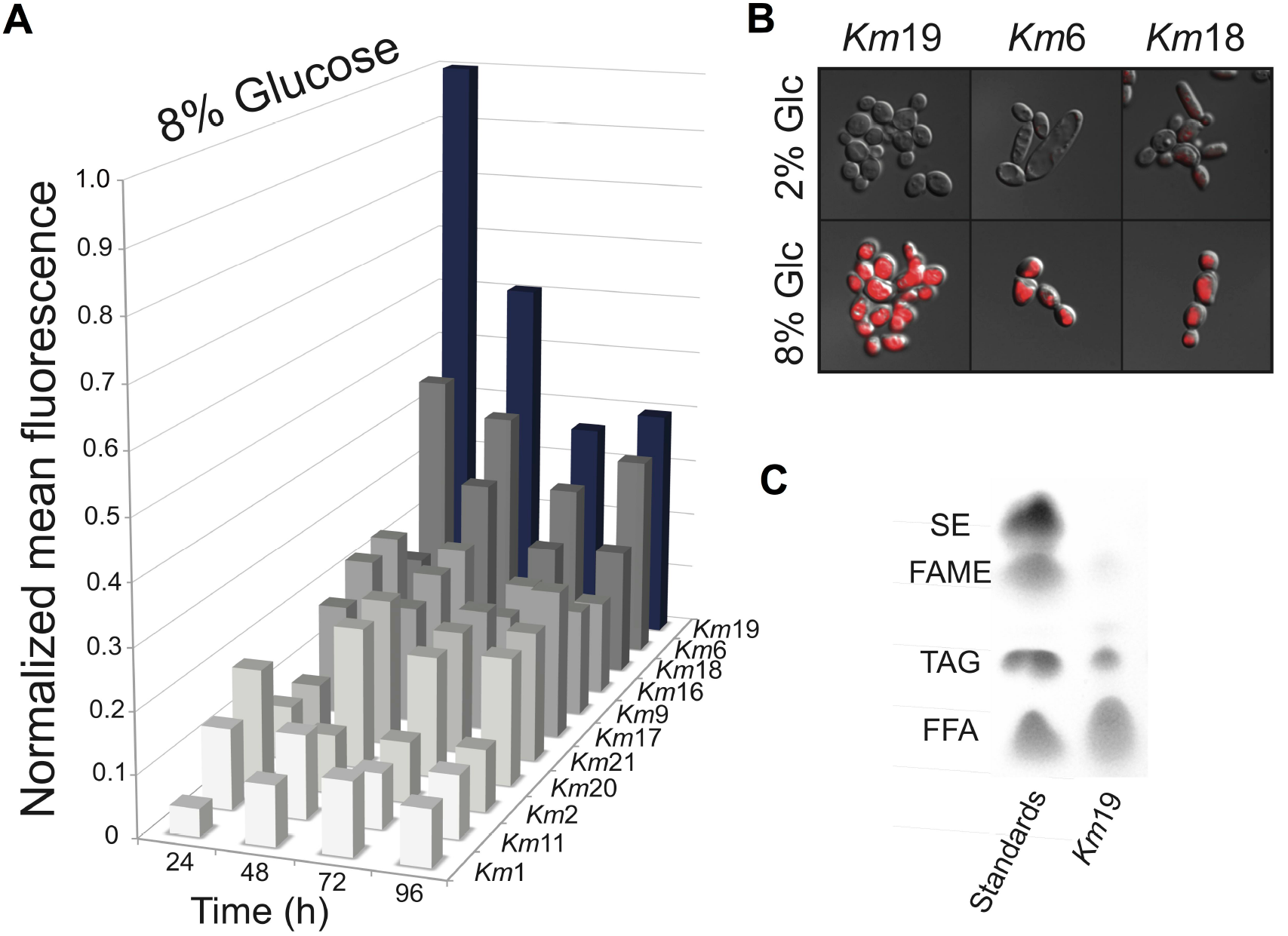
Lipogenesis of *K. marxianus* strains. **(A)** Nile red fluorescence flow cytometry of 11 wild type isolates after 24, 48, 72 and 96 hours at 42 °C in lipogenesis medium. **(B)** DIC images superimposed with epifluorescence microscopy of Nile red stained cells. Little or no fluorescence is seen after 24 hours in 2% glucose. After 24 hours in 8% glucose at 42 °C (*Km*19 and *Km*6) and 48 hours (*Km*18) fluorescence is seen encompassing the majority of the cell volume. **(C)** TLC analysis of *Km*19 total lipids after 24 hours in 8% glucose at 42 °C. Lane 1, standards ladder containing SE, steryl ester; FAME, fatty acid methyl ester; TAG, triacylglycerols; FFA, free fatty acids. Lane 2, *Km*19 lipids.

We used fluorescence microscopy to examine the cell morphology of the strains with the highest lipid titers. When Nile red fluorescence was overlaid with differential interference contrast images of *K. marxianus* isolates *Km*19, *Km*6, and *Km*18 (**Table 1**) after 24 or 48 hours of growth in 8% glucose at 42 °C, large lipid droplets encompassed a large fraction of the cell volume (**Fig. 3*B***). *Km*19 produced the highest levels of lipids as measured by Nile red fluorescence, which peaked after only 24 hours (**Fig. 3*A***), at which point *Km*19 had accumulated lipids at ~10% dry cell weight (*SI Appendix*, Fig. S6). Thin layer chromatography (TLC) revealed that the majority of the lipid in *Km*19 accumulated as free fatty acids (FFA) (**Fig. 3*C***).

### Strain engineering for higher lipid production

Lipogenesis in oleaginous yeasts such as *Yarrowia lipolytica* results in the synthesis and storage of lipid droplets within the cytoplasm (22). Lipid biosynthesis is largely dependent upon the enzymes AMP deaminase (AMPD), ATP-citrate lyase (ACL), acetyl-CoA carboxylase (ACC), and malic enzyme (MAE) (23). Collectively, these enzymes promote the accumulation of acetyl-CoA via citrate. Interestingly, although ACL is thought to be crucial for lipogenesis in oleaginous yeasts (24), we did not identify the genes *ACL1* and *ACL2* in the *K. marxianus* reference strain *Km*1. ACC1 then converts acetyl-CoA into malonyl-CoA and the malic enzyme provides NADPH, the reduced cofactor necessary for the production of lipids. Total lipid accumulation is a balance between lipid synthesis and catabolism through β-oxidation in the peroxisome (**Fig. 4*A***). Engineered strains of *Y. lipolytica* with reduced β-oxidation (*pex10Δ*) and peroxisome biogenesis (*mfe1Δ*) combined with over-expression of lipogenesis enzymes can store up to 80-90% dry cell weight as lipid compared to only ~10-15% lipid content for wild type cells (23). However, the high yields of lipogenesis in *Y. lipolytica* often takes up to 5 days to reach its peak (25), and requires temperatures of ~30 °C, due to the lack of thermotolerance in this yeast (23).

**Figure 4.**
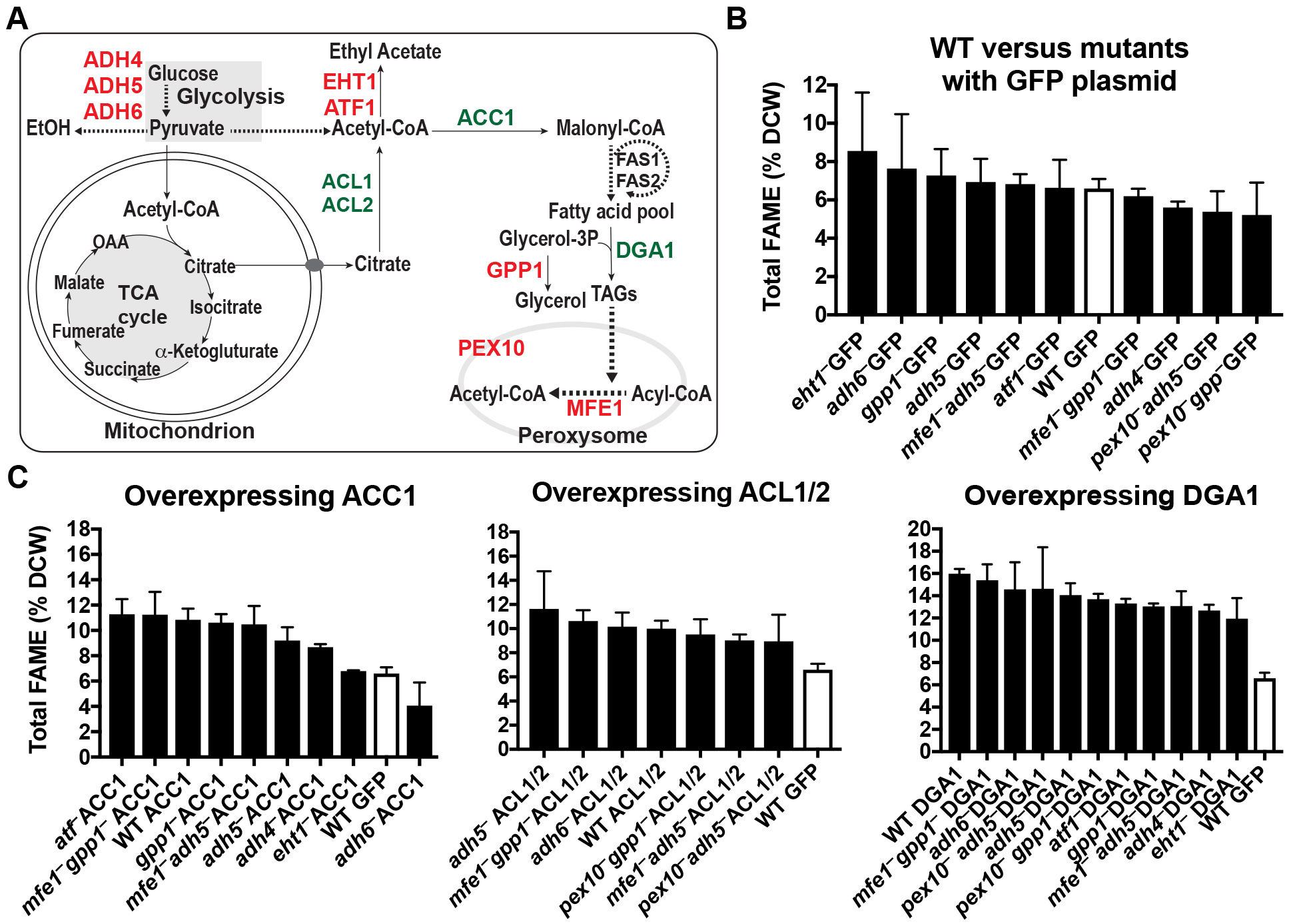
Genetic dissection of lipogenesis of a high-producing *K. marxianus* strain, *Km*6. **(A)** General overview of lipid-related metabolism. Genes in red were inactivated with CRISPR_NHEJ_ and genes in green were overexpressed using plasmids. (**B**) Percentage of fatty acids in dry cell weight after 24 hours in lipogenic conditions at 42 °C for several variants of *Km*6 (wild type and mutants). (**C**) Percentage of fatty acids in dry cell weight for several *Km*6 variants containing *ACC1, DGA1* and *ACL1/2* overexpression plasmids. In panels (**B**) and (**C**) all experiments were carried out in biological triplicate, with mean values and standard deviations shown. Lipogenesis medium in (**B**) and (**C**) contained monosodium glutamate instead of ammonium sulfate.

We tested whether inactivation or over-expression of genes previously shown in *Y. lipolytica* to contribute to high lipid production (23) would affect the ability of *K. marxianus* to produce lipids. Although wild-type *Km*19 produced the most lipids of the wild-type strains we tested, *Km*19 is very difficult to transform with plasmids. Therefore, we chose to use strain *Km*6 (**Table 1**), since it has similar lipid content and is easily transformed with plasmid DNA, allowing the facile use of plasmid-based CRISPR-Cas9 to inactivate genes, or plasmid-based over-expression.

Unlike *Y. lipolytica*, fermentation of glucose to ethanol and esterification of acetate to ethyl acetate are likely to compete with lipogenesis in *K. marxianus*. Therefore, we inactivated genes by CRISPR_NHEJ_ to decrease ethanol fermentation (*ADH* genes) (26), and ethyl acetate production (*ATF* genes) (7) as well as ester biosynthesis *(EHT1)* (7, 27) and glycerol biosynthesis *(GPP1).* We also inactivated genes involved in β-oxidation (*PEX10* and *MFE1*) (23) (**Fig. 4*A***). For the overexpression experiments, we cloned genes known to be involved in the accumulation of lipids *(DGA1, ACC1* and the dimer *ACL1/ACL2*) into overexpression plasmids constructed using strong promoters (*KmTDH3* or *KmPGK1*) to drive the expression of these genes. The plasmids were individually transformed into wild type *Km*6 or strains in which CRISPR_NHEJ_ had been used for targeted gene inactivation. The fatty acid content of each of these engineered strains was measured by gas chromatography and calculated as the percentage of dry cell weight. Although none of the inactivated genes had an appreciable effect in the accumulation of fatty acids (**Fig. 4*B***), over-expressing *DGA1* increased the levels of fatty acids across all strains tested, more than doubling the fatty acid content in the wild-type strain (**Fig. 4*C***). No appreciable differences were found in terms of the fatty acid composition except for the *Km*6 *eht1*^−^ strain bearing the *ACC1* plasmid which had more than 80% of its total fatty acids comprised of stearic acid (18:0), compared to ~10-20% in the other strains (*SI Appendix*, Fig. S7).

### Breeding to isolate high-producing, thermotolerant, and transformable strains

The ability to cross phenotypically diverse and stable haploid *K. marxianus* strains should enable combining several beneficial traits into a single strain. For example, strain *Km*19 is the best lipid-producing strain we identified (**Fig. 3*A***), but it is neither easily transformed nor thermotolerant when compared to other *K. marxianus* strains. On the other hand, strain *Km*17 transforms easily, is thermotolerant (growing at 45 °C), but is only moderately oleaginous (**Fig. 3*A***). We therefore crossed these two strains to combine their beneficial traits into single isolates. We first engineered *Km*19 α*3*^−^ *kat1*^−^ *trp1*^−^ using CRISPR_NHEJ_ and crossed a *MAT***a** isolate with an engineered stable haploid *Km*17 *MAT*α α*3*^−^ *kat1*^−^ *leu2*^−^ strain. The resulting diploids were sporulated, then germinated at high temperature (44 °C) to select for thermotolerant segregants, and 91 haploid progeny were picked to screen for lipid production and plasmid transformability (**Fig. 5*A***).

**Figure 5.**
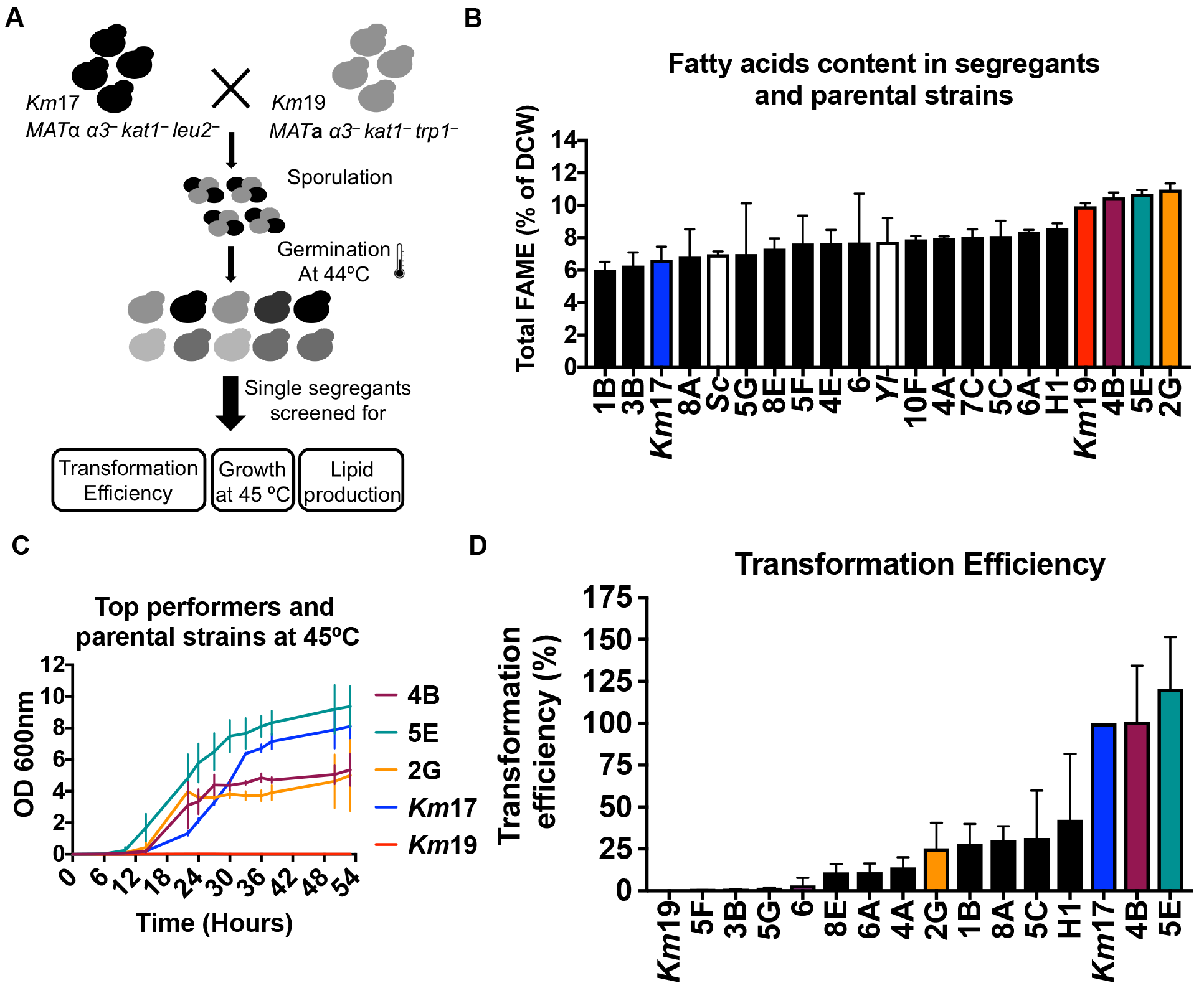
Selection of *K. marxianus* strains with combined beneficial traits. (**A**) Selection strategy. *Km*19 and *Km*17 were crossed, sporulated and then germinated at 44 °C to select for thermotolerant segregants. Single segregants were isolated and tested for lipid production, transformability and high temperature growth. (**B**) Fatty acid percentage in dry cell weight for several segregants from the *Km*17 × *Km*19 cross and the parental strains. Three segregants have similar profiles to the more lipogenic parental strain (*Km*19). Experiments are from biological triplicates with mean and standard deviation shown. (**C**) Growth curves at 45 °C for the segregants 4B, 5E and 2G as well as parental strains *Km*17 and *Km*19, in biological triplicate. *Km*19 is unable to grow at this temperature. Growth curves for parental strains at 30 °C, 37 °C and 42 °C can be found in the supporting information (*SI Appendix*, Fig. S9) as well as for segregants at 30 °C and 37 °C (*SI Appendix*, Fig. S10). (**D**) Transformation efficiency for several segregants normalized by *Km*17 transformation efficiency. Experiments are from 2-4 biological replicates with normalized mean and standard deviations shown.

Single segregants isolated from the above temperature selection were individually scored in terms of lipid production using Nile red staining and flow cytometry, and displayed high variability in lipid production (*SI Appendix*, Fig. S8). A few segregants performed better than the parental *Km*19 haploid strain, but a number of these had high fluorescence due to aggregation as determined by light microscopy, and were therefore excluded from further analysis. Strains that did not aggregate had their fatty acid percentage in dry cell weight and composition measured using GC-FID (**Fig. 5*B***). Notably, three of these isolates produced lipids as well as the parent *Km*19 strain, while inheriting the thermotolerance and transformability of *Km*17 (**Fig. 5*C*, 5*D***, SI Appendix Fig. S9, S10). These strains therefore combined all three beneficial traits of the parental strains.

## DISCUSSION

Common metabolic engineering techniques are not ideal when dealing with complex phenotypes such as thermotolerance, productivity and robustness (28). Therefore, agnostic approaches for combining complex traits into model yeast species are of high value, including directed evolution (29), genome-wide transcriptome engineering (30), and genome shuffling (31). However, these cannot substitute for classical sexual crossing for combining strain-specific traits. It is known that sexual reproduction enables adaptation to stressful industrial environments due to the faster unlinking of deleterious allelic pairs compared to clonal populations (10). Here we establish *K. marxianus* as a platform for synthetic biology by engineering stable heterothallic haploid strains that can be crossed to combine complex, unmapped multigenic traits into one strain.

In *S. cerevisiae*, inactivating a single gene encoding HO endonuclease makes this yeast heterothallic, and is sufficient to gain laboratory control of its mating cycle (32). While inactivating the single transposase α3 can be used to cross *K. marxianus* strains (8), the resulting strains possess unstable mating types due to the presence of Kat1, and could randomly switch from *MAT***a** to *MAT*α (**Fig. 1*B***, **Fig. 2*C***, *SI Appendix*, Fig. S4). To establish stable crossing in *K. marxianus* and abolish self-mating, we used CRISPR_NHEJ_ to inactivate both transposases that are responsible for mating type switching (α3 and *Kat1*) (**Fig. 1*B***). By simultaneously inactivating *ALPHA3* and *KAT1*, we created stable heterothallic α*3*^−^ *kat1*^−^ strains which cannot switch mating type and, therefore, can be mated in a controlled manner with strains of the opposite mating type (**Fig. 2*A***, *SI Appendix*, Fig. S4). Alternative methods for creating stable haploids by deleting the silenced *MAT* loci have been used in yeast (33) but this strategy creates sterile strains and can be lethal if the endonuclease that initiates the double strand break required for mating type switching is not inactivated as well (34). Our strategy of inactivating both the α3 and Kat1 transposases preserves the ability to mate *K. marxianus* strains, an essential tool for synthetic biology, and to take advantage of this yeast’s remarkable phenotypic diversity. We successfully isolated heterothallic haploid strains from 12 wild type isolates. These strains readily mate with each other, resulting in sporulation-competent diploids that segregate to viable haploid spores. Combined with CRISPR-Cas9 genome editing (**Fig. 1**) (6–9), these results establish a full set of tools for use of *K. marxianus* as a synthetic biology host and future exploration of its biology on a genome-wide scale.

Using both sets of synthetic biology tools described here, we sought to exploit the diversity of *K. marxianus* as a thermotolerant, fast growing, Crabtree-negative yeast. Screening 11 of the wild-type isolates for high levels of lipogenesis, we found high strain-to-strain variability in lipid production. Notably, strain *Km*19 produced ~10% lipid by dry cell weight after 24 hours at 42 °C, a considerably shorter time than the 120 hours required by wild-type *Y. lipolytica* to accumulate a similar amount of lipid (25). We find *Km*19 stores the vast majority of lipid as free fatty acids (FFAs) (**Fig. 3*C***), in contrast to *Y. lipolytica*, which stores lipids as triacylglycerols (22, 35). FFAs are particularly suitable for the production of alkanes/alkenes and fatty alcohols, two types of high-value chemicals (36, 37). Although *Km*19 produces lipids at a rapid rate, it is not thermotolerant compared to other *K. marxianus* isolates (**Fig. 5*C***), and is difficult to transform with plasmid DNA. To eliminate these barriers to conducting genetic engineering in the oleaginous strain *Km*19, we crossed it with *Km*17, which transforms well and grows well at 45 °C. Notably, some progeny of this cross isolated as single segregants produced lipids to the same level of the parent *Km*19 strain while retaining thermotolerance and transformability of *Km*17, and were also not prone to aggregation. The power of combining multiple complex and valuable traits using stable heterothallic strains, together with CRISPR-Cas9 genome engineering, opens a new frontier to use *K. marxianus* as both a thermotolerant model species and an industrially-relevant host.

## MATERIALS AND METHODS

A full description of the materials and methods used in this study, including media preparation, plasmid construction, high efficiency DNA transformation, CRISPR_NHEJ_ and CRISPR_HDR_ protocols, auxotrophic mating assays, sporulation and spore purification and determination of fatty acids by gravimetric analysis and GC-FID is provided in the *SI Appendix*.

## ACKNOWLEDGMENTS

We thank Jasper Rine, Jacob Corn and Christopher Somerville for valuable discussion and input during the course of this research. This work was funded by the Energy Biosciences Institute (J.H.D.C., J.W.T., and A.P.A.), the National Science Foundation (MCB-1244557 to J.A. Doudna), the CAPES Science Without Borders program (fellowship 18813127 to R.E.), FAPESP (fellowship 2017/24957-3 to M.F.) and the Bakar Spark Fund (J.H.D.C.).

## SUPPORTING INFORMATION

### SI MATERIALS AND METHODS

#### Strains, media and culture conditions

*K. marxianus* strains used in this study were purchased from ATCC (American Type Culture Collection), CBS (The Dutch Centraalbureau voor Schimmelcultures, Fungal Biodiversity Centre) or were obtained from an in-house collection. A complete list of all the wild-type strains is given in **Table 1**. Strains *Km*1, *Km*20, and *Km*21 have internal identity codes YST31, 1S300000, and 1S1600000, respectively. Strains were stored at −80 °C in 25% glycerol. All experiments began by inoculating a 12 mL culture tube containing 3 mL yeast peptone dextrose (YPD) or Synthetic Complete Dextrose (SCD) media with a single colony grown on YPD media agar plate. Cultures were shaken at 250 RPM. YPD agar consisted of 10 g/L yeast extract, 20 g/L peptone, 20 g/L glucose and 20g/L agar. SCD consisted of 2 g/L Yeast Nitrogen Base without amino acids or ammonium sulphate, 1 g/L CSM, 5 g/L (NH_4_)_2_SO_4_. 5% Malt extract media was made by mixing 30 g of malt extract with 20 g of agar and bringing the volume to 1 L with H_2_O, then sterilized by autoclaving at 10 psi for 15 minutes. Sporulation medium (SPO) was made with 10 g/L potassium acetate, 1 g/L bacto-yeast extract, 0.5 g/L glucose. 5-FOA plates contained 2 g/L Yeast Nitrogen Base without amino acids or ammonium sulphate, 5 g/L (NH_4_)_2_SO_4_, 1 g/L complete CSM, 20 g/L glucose, 20 g/L agar, and 1 g/L 5-Fluoroorotic Acid (5-FOA). Lipogenesis medium contained 2g/L YNB without amino acids and ammonium sulfate, 1 g/L ammonium sulfate or 1 g/L monossodium glutamate and 8% glucose or cellobiose.

#### Genome sequencing and annotation

A single colony isolate of strain YST31 (**Table 1**, *Km*1) grown on a YPD plate was used to inoculate a YPD liquid culture and prepare genomic DNA using the YeaStar™ Genomic DNA Kit (Zymo Research). We submitted ~5 mg of genomic DNA for small insert library preparation (~250 bp) and Illumina sequencing. Library preparation and genome sequencing (Illumina HiSeq 2500) was performed by the UC Davis Genome Center DNA technologies core (http://dnatech.genomecenter.ucdavis.edu/). For YST31, we obtained 14,790,917 PE100 paired-end reads and after trimming we assembled reads into 116 scaffolds using CLC Genomics Workbench version 7.5.1. Default settings were used for quality trimming and *de novo* assembly. The median coverage was 250-fold and the total genome assembly was 10,784,526 bp. Genome annotation was performed using an automated software pipeline FGENESH++ (http://www.softberry.com) version 3.1.1. Genes were first predicted *ab initio* using FGENESH and then refined based on protein homology (38, 39). A custom BLAST database based on Genbank nr (downloaded Oct. 31, 2014) was used for homology refinement of gene models. Gene prediction parameters were obtained from Softberry and were based on *Saccharomyces cerevisiae* gene models as the training set. The resulting annotation output files were renumbered and converted into Genbank format using custom scripts provided by Softberry. The raw fastq reads have been deposited in the NCBI SRA and the scaffolds and annotation in Genbank with accession numbers <u>X</u> and <u>Y</u>, respectively.

#### Cas9 plasmid construction

To manipulate *Kluyveromyces marxianus*, we created a plasmid that can replicate in both *Escherichia coli* and *K. marxianus* and confer resistance to kanamycin and Geneticin, respectively. We used plasmid pOR1.1 (13), which can replicate in *E. coli* and *S. cerevisiae*, as a backbone for further manipulation. We identified and cloned an autonomous replicating sequence (ARS) from commercially available *K. marxianus* strain ATCC 36907 (*Km*11, **Table 1**) as follows. Using a YeaStar genomic DNA extraction kit (Zymo Research, D2002), genomic DNA was extracted from *K. marxianus* ATCC 36907. One microgram of genomic DNA was incubated with restriction enzyme EcoRI (NEB, R0101S) to fragment the DNA. In parallel, the *S. cerevisiae* 2μ origin of replication was replaced with an EcoRI digestion site in pOR1.1. The plasmid was then linearized with EcoRI and treated with Shrimp Alkaline Phosphatase (Affymetrix, 78390) to dephosphorylate the DNA ends and prevent re-ligation of the vector. The genomic DNA fragment pool was ligated with the linearized plasmid using T4 DNA Ligase (Invitrogen, 15224017), transformed into One Shot TOP10 competent *E. coli* (C404003), and plated on kanamycin selection plates. All growing colonies were pooled, and the plasmids extracted using QIAprep Spin Miniprep Kit (Qiagen, 27106). Two micrograms of the resultant plasmid pool were transformed into ATCC 36907 and plated on Geneticin selection plates. Many colonies were picked, and plasmid extraction was performed for each using the Zymo Research Yeast Plasmid Extraction Kit (D2001). The plasmids were individually transformed back into One Shot TOP10 competent *E. coli*, the plasmids extracted once more, and digested with EcoRI. The digests were run on a 1% agarose gel with TAE buffer (40), and the clone with the smallest insert was chosen. The insert was sequenced and then systematically trimmed to a 232 bp functional region that still conferred the ability of the plasmid to replicate in *K. marxianus* (*SI Appendix*, **Table S3**).

The resulting plasmid with a *K. marxianus* ARS was then modified to express Cas9 and a single-guide RNA (sgRNA) cassette using transcription promoters and terminators from *K. marxianus.* The *S. cerevisiae* promoter and terminator for Cas9 as used in pCas (13) were replaced with those for homologous genes in *K. marxianus.* In the new plasmid, Cas9 expression was driven by the promoter region of the gene *KmRNR2* - a mild-strength promoter - and terminated by the strong *KmCYC1* terminator. *S. cerevisiae* tRNAs were used as promoters to drive sgRNA expression (13) and terminated by the *ScSNR52* terminator. Between the promoter and the sgRNA there is a hepatitis δ-ribozyme sequence that cleaves off the 5′ leader sequence, liberating the tRNA from the sgRNA body that binds to Cas9 protein. The released transcript contains the δ-ribozyme, the protospacer sequence that targets Cas9 to the desired sequence and the scaffold sgRNA (13).

#### Overexpression plasmid construction

Four genes found to be involved in lipogenesis in other yeast were cloned into overexpression plasmid backbones using In-Fusion Cloning Kit (Takara). The cloning reactions contained 25-50 ng of vector, 3 times molar excess of PCR-generated insert and 0.5 μL of In-Fusion in a final volume of 2.5 μL. *K. marxianus ACC1* and *DGA1* coding sequences were amplified from *Km*6 gDNA using Phusion polymerase and cloned into two different linearized backbones, while *Yarrowia lipolytica ACL1* and *ACL2* coding sequences were cloned into the same plasmid. The vector backbone was the same used for pKCas9 construction, containing a *K. marxianus* ARS isolated as described above, a Geneticin resistance marker and pUC bacterial origin of replication. *ACC1* and *ACL1* were controlled by the *K. marxianus GK1* promoter, while *DGA1* and ACL2 were under control of the *TDH3* promoter (*SI Appendix*, Table S3). All ORFs were terminated by the *K. marxianus CYC1* terminator sequence. The resulting plasmids were transformed into wild type *K. marxianus* strain *Km*6 (**Table 1**) as well as into 10 knockout mutant strains derived from *Km*6 that were constructed using CRISPR_NHEJ_: *adh5*^−^, *adh6*^−^, *adh4*^−^, *gpp1*^−^, *atf1*^−^, *eht1*^−^, *mfe1*^−^ *gpp1*^−^, *mfe1*^−^ *adh5*^−^, *pex10*^−^ *gpp1*^−^, *pex10*^−^ *adh5*^−^ (*SI Appendix*, Table S1.

#### High efficiency DNA transformation

We established a high-efficiency transformation protocol for *K. marxianus* as follows. A single colony was inoculated in 1.5 mL of YPD media and incubated at 30 °C overnight, then 180 μL of this culture was transferred to 5 mL of fresh YPD media and incubated at 30 °C until the OD_600_ reached 1.0-1.2 (~5 to 6 hours). Then, 1.4 mL of the culture was aliquoted into microcentrifuge tubes and spun down at 3,000 g for 5 minutes. The supernatant was removed and the pellet resuspended in 50 mM of lithium acetate, followed by incubation at room temperature for 15 minutes. The cells were spun down, the supernatant discarded and the cells used for subsequent transformation reactions.

Single-stranded DNA (ssDNA) was previously prepared as follows (41): 2 μg/μL of ssDNA from Sigma (D1626-250mg) was agitated with a stir bar overnight in TE buffer (10 mM Tris, pH 8.0 and 1 mM EDTA) at 4 °C, then concentrated to 10 μg/μL by isopropanol precipitation, resuspended in water and quantified (NanoDrop™ 1000, Thermo Scientific). Prior to each transformation, aliquots of ssDNA were boiled for 5 minutes then placed in ice bath for 5 minutes. Keeping all the reagents on ice, 66.7 μL of 60% PEG 2,050, 12.5 μL of 2 M lithium acetate, and water in a final volume of 100 μL were added to a sterile microcentrifuge tube. Then 2 μL of 1 M DTT and 25 μg of ssDNA were added, followed by 0.1 μg to 5 μg of pKCas plasmid DNA. The transformation mix was briefly vortexed and 100 μL was added to the cells. The transformation reaction was then incubated at 42 °C for 40 minutes. The reaction was spun down for 5 minutes at 3,000 g, the supernatant removed and fresh 500 μL of YPD added. The cells were allowed to recover for 2 hours at 37 °C and 250 RPM. Then 10% of the volume was spread on a YPD G418 selection plate, and the remaining volume spread on a second G418 plate.

#### CRISPR_NHEJ_ and CRISPR_HDR_

We transformed the pKCas plasmid into *K. marxianus* strains in the absence of donor repair DNA to determine the efficiency of NHEJ repair of the double strand break. We cloned the guide sequence for the sgRNA in pKCas to target *URA3 (SI Appendix*, Table S3), to allow counter-selection with 5-fluoroorotic acid (5-FOA) plates, which select for *ura3*^−^ colonies. Approximately 1 μg of pKCas plasmid was transformed into *K. marxianus* and the efficiency of editing was calculated by counting the number of *ura3~* colonies divided by the number of G418^R^ transformants. Sequencing of the targeted region revealed small insertions or deletions (indels) around the Cas9 cleavage site typical of NHEJ, resulting in premature stop codons within the *URA3* ORF (*SI Appendix*, Fig. S1). We find that this system can be used to create inactive alleles in different genes with efficiencies of ~75% (*SI Appendix*, Fig. S1).

We modified the high-efficiency transformation protocol to enable cotransformation of the pKCas plasmid and a linear repair DNA template (donor DNA), for HDR-mediated genome editing. For tests of HDR-mediated gene insertion, the donor DNA targeted for genome integration contained the NatMX cassette conferring resistance to Nourseothricin and 0.9 kb of flanking homology to the target site in the *K. marxianus* genome. Donor DNA was generated by PCR and concentrated by isopropanol precipitation (42). The best ratio of plasmid to donor DNA was 0.2 μg of pKCas plasmid and 5 μg of linear donor DNA, with 0.9 kb of homology to the Cas9 targeting site. Tests with higher concentrations of both pKCas and donor DNA were not as efficient. The transformation reaction was carried out as described above, except that cells were allowed to recover for 1 hour at 37 °C in YPD without drug at 250 RPM, after which G418 was added and the cells were allowed to recover overnight. For the Nourseothricin gene insertion experiments, cells were plated on YPD G418 plates, incubated at 37 °C overnight and then replica-plated on either 5-FOA or Nourseothricin plates, to identify Nat^R^ colonies with correct insertion in the *URA3* locus. Colony PCR performed on select Nat^R^ colonies (targeting outside the 0.9 kb homology arms of the NatMX cassette) confirmed NatMX cassette integration at the *URA3* locus.

#### Mating-competent heterothallic strains

*ALPHA3* and *KAT1* double inactivation strains (*α3*^−^ *kat1*^−^) were constructed using CRISPR_nhej_ (*SI Appendix*, Table S1). *K. marxianus* was transformed with either *KAT1* or *ALPHA3* targeting pKCas plasmids. Genomic DNA was isolated from G418^R^ colonies, the *KAT1* or *ALPHA3* regions were PCR-amplified and the PCR products sequenced. Colonies with sequences containing indels that generated early stop codons were chosen and were saved as glycerol stocks. Single inactivation strains were subjected to a second round of CRISPR_NHEJ_ to inactivate the second transposase, creating the double mutants *α3*^−^ *kat1*^−^. These double mutants were then subjected to a third round of CRISPR_NHEJ_ targeting the *LEU2* or *TRP1* genes to create auxotrophic strains. The resulting strains were tested for mating type and heterothallic status by using a pheromone assay described below, or by crossing them with reference heterothallic haploid strains, allowing haploid heterothaliic *MAT***a** or *MAT*α strains to be successfully isolated. Some double-transposase inactivated strains did not mate with the reference strains, possibly because these are stable diploids or triploids (12) or possibly due to chromosomal rearrangements (43). Although we performed the gene inactivations individually, we later tested and verified that simultaneous targeting of both transposases with CRISPR_NHEJ_ works efficiently in *K. marxianus*.

#### Mating pheromone response assay

We used reciprocal BLASTp using *S. cerevisiae* and *K. lactis* protein sequences as queries to identify two putative *MFA* genes (*KmMFA1* and *KmMFA2*) as well as the *MFμ* gene (16, 17). The sequences in *S. cerevisiae* are encoded by genes YPL187W and YGL089C in the Saccharomyces Genome Database (44) and those in *K. lactis* are encoded by Genbank entry CAG99901.1 (REFSEQ ID XP_454814.1) and as described in (17). To the best of our knowledge, these pheromones had not been previously identified in *K. marxianus.* We found two putative *MFA* genes (*KmMFA1* and *KmMFA2*) as well as the *MF*α gene (*Km*MFα1 which encodes 2 isotypes *Km*MFα1 and *Km*MFα2) by reciprocal BLASTp using the *S. cerevisiae* and *K. lactis* protein sequences as queries (*SI Appendix*, Table S2). Interestingly, the deduced **a**-factor amino acid sequences of *Km*MFa1 and *Km*MFa2 are not completely conserved, differing by 1 amino acid (**Fig. 1*C***). Sequencing *KmMFA1*, *KmMFA2* and *KmMF*α from 8 unique strains show full strain-to-strain conservation. Notably, *Km*MFa1 is completely conserved with the respective *K. lactis* sequence, suggesting a relatively conserved sexual cycle.

Synthetic **a**- and α-pheromones were obtained from Genemed Synthesis Inc. and resuspended in water. To verify if the putative α-factors (*Km*MFα1 and *Km*MFα2) are viable and induce mating projections (“shmoo”) in engineered heterothallic cells, we performed a pheromone morphological response assay. Cells were grown in YPD medium to a density of 10^6^ cells/100 mL and both pheromones were added to a final concentration of 25 mg/mL. Cells were then examined under the light microscope at different times. Mating projections were observed after 6 hours for the mature α-factors, *Km*Mfα1 and *Km*MFα2, while no morphological differences were seen in the absence of synthetic pheromone. Strains sensitive to the α-pheromones were classified as putative *MAT***a**, and strains that exhibited no mating projections in the presence of either α pheromone for 12 hours were presumed to be stable *MAT*α haploids, or *MAT***a**/*MAT*α diploid strains. The mating type of heterothallic strains was verified by sequencing of the *MAT* loci, by PCR amplification of the *MAT* loci with primers flanking the *MAT* loci, and *MAT* specific primers (*SI Appendix*, Table S2 and Fig. S7). The same procedure was done using the synthesized **a**-factors but they failed to induce mating projections in the strains tested, possibly because **a**-factors are reported to be heavily post-translationally modified unlike α-factors (45).

#### *K. marxianus* auxotrophic mating

Auxotrophic double mutant strains (α*3*^−^ *kat1*^−^) of either mating type were grown up overnight from a single colony in 5 mL SCD media. Strains were pelleted at 3,500 g for 5 minutes followed by washing with 1 mL of sterile water. Strains were pelleted again and resuspended in 50 μL of water. 4 μL of the cell suspension was dispensed as a single drop (patched) onto 2% glucose plates or MA5 plates and allowed to dry. Strains to be mated containing a complementary auxotrophic marker (either *leu2*^−^ or *trp1*^−^) were dispensed on top of previous dried spots and allowed to mate at room temperature for 24-48 hours. Mating plates were replica-plated onto SCD agar plates minus both leucine and tryptophan. Diploids cells were grown at 30 °C for 48 hours. Freezer stocks were made by scraping off the diploid patch, resuspending in 25% glycerol, and freezing at −80 °C.

#### Sporulation and spore purification

For sporulation, diploid strains were taken from freezer stocks and streaked onto an SCD agar plate minus leucine and tryptophan to ensure diploid strain growth. Single colonies were inoculated in 5 mL SCD liquid media and grown overnight at 30 °C. Many strains (including *Km*17) were observed to be unstable as diploids and this treatment alone resulted in 25-90% sporulation. Strains or crosses that resulted in relatively stable diploids, were then patched onto MA5 or SPO plates or suspended in 1 mL of M5 or SPO liquid media and incubated at room temperature. Cells were observed under the optical microscope every 24 hours for sporulation. Typically, sporulation was rapid and resulted in 25-95% sporulation in 24 hours. However, recalcitrant strains would require up to 3-4 days.

Spore purification was performed as described previously (46). Diploid cultures from sporulation media (MA5 or SPO media) were scraped from solid media or centrifuged at 3,500 g from liquid media for 5 minutes, re-suspended into softening buffer (10 mM DTT, 100 mM Tris-SO_4_, pH 9.4) at a cell/spore density of ~1 × 10^8^ /mL and incubated for 10 minutes at 30 °C. The spore/cell suspensions were centrifuged at 3.500 g and suspended in spheroplasting buffer (2.1 M sorbitol, 10 mM potassium phosphate, pH 7.2) to ~3 × 10^8^ cell-spores/mL. Zymolyase-100T (United States Biological Corporation; Fisher Scientific) was added to a final concentration of 0.2 mg/mL and incubated at 30 °C for 20 min. Spheroplasts and spores were centrifuged at 3.500 g for 5 minutes and spheroplasts were lysed by multiple washes with 0.5% Triton X-100.

#### Complementation mating assay

To further validate the role of α3 and Kat1 in switching mating types, we constructed expression plasmids containing the coding sequences of each protein to perform a complementation assay. The proteins were cloned into the same vector used for Cas9 expression, under control of the moderate-strength promoter of the gene *KmRNR2.* Alternative plasmids bearing the native promoter of *KAT1* or *ALPHA3* were also constructed. These plasmids were transformed into *α3*^−^ *kat1*^−^ *leu2*^−^strains of either mating type. To test transformants for mating-type switching, we crossed them with stable heterothallic reference strains of either *MAT***a** or *MAT*α mating types that also lack the ability to produce tryptophan (*α3*^−^ *kat1*^−^ *trp1*^−^). We inoculated 12 single transformats of each mating type in 20 mL of SCD medium and incubated for 24 hours at 30 °C and 220 RPM shaking. The cultures were then washed and resuspended in 70 μL of water. 4 μL of each cell concentrate were spotted onto glucose 2% agar plates to induce switching and, later, mating. Reference strains of the same and opposite mating type were spotted on top of the 12 spots of transformants. The plates were incubated for 24 hours or 48 hours at room temperature and then replica-plated onto SCD minus (Leu, Trp) plates. Growth was only observed in spots where mating occurred.

#### Culture conditions for inducing lipogenesis

Single colonies were grown overnight in 3 mL of YPD medium at 30 °C and 225 RPM. Then 20 mL of YPD media was inoculated with 0.5 mL of the overnight culture and grown at 30 °C and 225 RPM for 24 hours. Cells were pelleted at 3,000 g for 10 minutes, washed with 1 mL of water and then transferred to 1.5 mL tubes before being pelleted again at 3,000 g for 10 minutes. The pellet was resuspended in 1 mL of lipogenesis media and transferred to a 12 mL round bottom tube containing 3 mL of the same media. For the strains containing overexpression plasmids, the lipogenesis medium had 1 g/L of monosodium glutamate instead of ammonium sulfate due to Geneticin incompatibility with ammonium sulfate. Cultures were then shaken at 275 RPM at 42 °C for 24 hours. Cells were harvested by transferring 250 μL of each culture to pre-weighed eppendorf tubes, spinning down at 3,000 g for 10 minutes, washing with 500 μL of water and then resuspending in 100 μL of water prior to freezing in liquid nitrogen. Cells were kept in the −80 °C prior to lyophilization. Culturing conditions for cellobiose-grown samples were similar except that lipogenesis media contained 8% cellobiose instead of glucose.

#### Epifluorescence microscopy and flow cytometry

To assay lipid production, *K. marxianus* cells were stained with the lipophilic dye Nile red (MP Biomedicals) which is permeable to yeast cells and a common indicator of intracellular lipid content. Single colonies of *K. marxianus* were grown in 3 mL of YPD media overnight for at 30 °C with 250 RPM shaking. Cell concentrations were then normalized, inoculated in fresh lipogenesis medium and grown for 24 hours at 42 °C at 250 RPM. Cells were harvested by spinning down 1 OD_600_ unit at 3,000 g for 3 minutes and then resuspended in 500 μL of PBS buffer (Phosphate-Buffered Saline solution, Sigma Aldrich). Then 6 μL of 1 mM Nile red solution (in DMSO) was added to the cells and incubated in the dark at room temperature for 15 minutes. Cells were spun down and washed in 800 μL of ice cold water, spun down again and resuspended in another 800 μL ice cold water. Nile red stained cells were examined on a Zeiss Axioskop 2 epifluorescence microscope at 553 nm excitation and 636 nm emission.

For flow cytometry, 300 μL cell solution was diluted in 1 mL of ice cold water and tested in the BD Biosciences Fortessa X20 FACS using a 25,000 cell count, a forward scatter of 250, a side scatter of 250 and the 535LP and 585/42BP filters for fluorescence detection. Fluorescence data was analysed using FlowJo software (Tree Star Inc., Ashland, OR, USA) and mean fluorescence values were obtained.

#### Isolation of high-lipid producing strains

Strains *Km*19 and *Km*17 were made heterothallic by CRISPR_NHEJ_ by inactivating both *KAT1* and *ALPHA3* genes as well as *TRP1* or *LEU2* (*SI Appendix*, Table S1) to allow selection of diploids in SCD minus (Leu, Trp) medium. Although *Km*19 does not transform well, by using our high efficiency transformation protocol we were able to recover a few colonies that transformed with the pKCas9 plasmid and allowed isolation of *α3*^−^ *kat1*^−^ auxotrophic strains. *Km*17 had a much higher transformation success rate and triple inactivation strains were easily constructed. To select for thermotolerant and lipogenic strains, *Km*19 *MAT***a** α*3*^−^ *kat1*^−^ *trp1*^−^ was crossed with *Km*17 *MAT*α *α3*^−^ *kat1*^−^ *leu2*^−^ on MA5 media (SI *Appendix*, Table S1). Diploid cells were sporulated and a spore suspension was made. Spores were germinated in YPD liquid media overnight at 30 °C, plated onto YPD plates and incubated at 44 °C to select for thermotolerant strains. 91 colonies were then selected and subjected to lipogenesis conditions for 24 hours. The strains were ranked by Nile red mean fluorescence (*Appendix*, Figure S6), and those prone to flocculation as determined by light microscopy were eliminated from further analysis.

#### Determination of fatty acids by gravimetric analysis and GC-FID

To measure the percentage of fatty acids in dry cell weight we extracted lipids from lyophilized cells and performed gravimetric analysis. Briefly, 250 μL of culture was pelleted at 4,000 g for 10 minutes in pre-weighed 1.5 mL microcentrifuge tubes (Metter Toledo Excellence XS205DU balance). After the supernatant was removed, the cell pellet was suspended in 0.5 mL of water, and pelleted again at 3,000 g for 10 minutes, after which the pellet was resuspended in 100 μL of water. The samples were frozen and stored at −80 °C. Frozen samples were then lyophilized overnight and weighed to calculate dry cell weight.

To measure the percentage of fatty acids in the different strains, we extracted lipids from lyophilized cells prepared as described above and analysed them by gas chromatography. Fatty acids were extracted and transesterified into fatty acid methyl esters (FAMEs) with methanol in the presence of an acid catalyst. The dry cell pellet was transferred to 15 mL glass conical screw top centrifuge tubes and 1 mL of methanolic HCl (3 N concentration) with 2% chloroform was added to the pellet. To ensure complete transesterification, an additional 2 mL of methanolic HCl (3 N) plus 2% chloroform was added. Approximately 100 μg of an internal standard (methyl tridecanoate) with its exact mass recorded was prepared in methanol and was added to the tube. The tube was sealed with a Teflon-coated screw cap and heated at 85 °C for 1.5 hour with vortexing every 15 minutes. The mixture was then cooled to room temperature, and the resulting FAMEs were extracted by the addition of 1 mL of hexanes followed by 30 seconds of vortexing. An organic top layer was obtained by centrifugation of the sample at 3,000 g for 10 minutes. The top layer was carefully collected and transferred to a GC vial. 1 μL was injected in split mode (1:10) onto a SP2330 capillary column (30 m × 0.25 mm μ 0.2 μm, Supelco). An Agilent 7890A gas chromatograph equipped with a flame ionization detector (FID) was used for analysis with the following instrumental settings: Injector temperature 250 °C, carrier gas: helium at 1 mL/min, temperature program: 140 °C, 3 minutes isocratic, 10 °C/min to 220 °C, 40 °C/min to 240 °C, 2.5 minutes isocratic.

#### Total oil extraction from dry yeast cells for thin layer chromatography

Total yeast oil was extracted following the protocol described by Jordi Folch *et. al* (47) for analysis by thin layer chromatography. Approximately 20 mg dry cell and 500 mg silica beads were weighed into a 2 mL centrifuge tube. 1 mL MeOH was added to the tube, and it was vortexed. Then, the tube was put into an aluminum block and bead-beaten 4 times for 30 seconds with 30 seconds resting intervals in between. The contents were transferred into a conical 15 mL glass centrifuge tube, and 0.25 mL MeOH was used to rinse the residuals on the small centrifuge tube. 2.5 mL CHCl3 was added, and the tube was briefly vortexed. The tube was shaken for 1 hour at 235 RPM. 250 μL of CHCl_3_/MeOH (2:1) and 1 mL of MgCl_2 (aq)_ (0.034 %) were added into the mix, and the tube was shaken for 10 minutes. The solution was vortexed for 30 seconds, centrifuged at 3,000 RPM for 5 minutes and the upper aqueous layer was removed. The resulting organic layer was washed with 1 mL 2N KCl/methanol (4:1, v/v), vortexed and centrifuged in the same way. The aqueous upper layer was removed, and the resulting organic layer was introduced with the artificial upper phase (chloroform/methanol/water 3:48:47). The resulting mixture was vortexed and centrifuged, and the upper layer was aspirated. This step involving the artificial upper phase was repeated until the white layer at the interface completely disappeared.

#### Thin layer chromatography *analysis of lipid composition*

TLC plates (7 × 10 cm) were preheated in 120° C oven for at least 10 minutes. A piece of paper towel/filter paper (7 × 10 cm) was added into a 600 mL beaker for saturation, and solvent system 1 (SS1 - petroleum ether/Et_2_O/AcOH (70:30:2)) was added until solvent reached about 0.5 to 1 cm in height. The beaker was covered with aluminum foil and parafilm, and the resulting setup was left alone for at least 10 minutes. Compounds were spotted on the preheated TLC plate, and the plate ran in SS1 until the solvent front was half way up the plate. The plate was dried in room temperature for 15 minutes. The plate was run using solvent system 2 (SS2, petroleum ether/Et_2_O (98:2)) until the solvent front nearly reached the top of the plate. The resulting TLC plate was dried in fume hood for 30 minutes before being immersed in MnCl_2_ charring solution (0.63 g MnCl_2_ × 4H_2_O, 60 mL H_2_O, 60 mL MeOH, 4 mL concentrated H_2_SO_4_) for 10 sec. The stained plate was developed in 120° C oven for approximately 20 minutes or until dark spots were observed.

**Figure S1.**
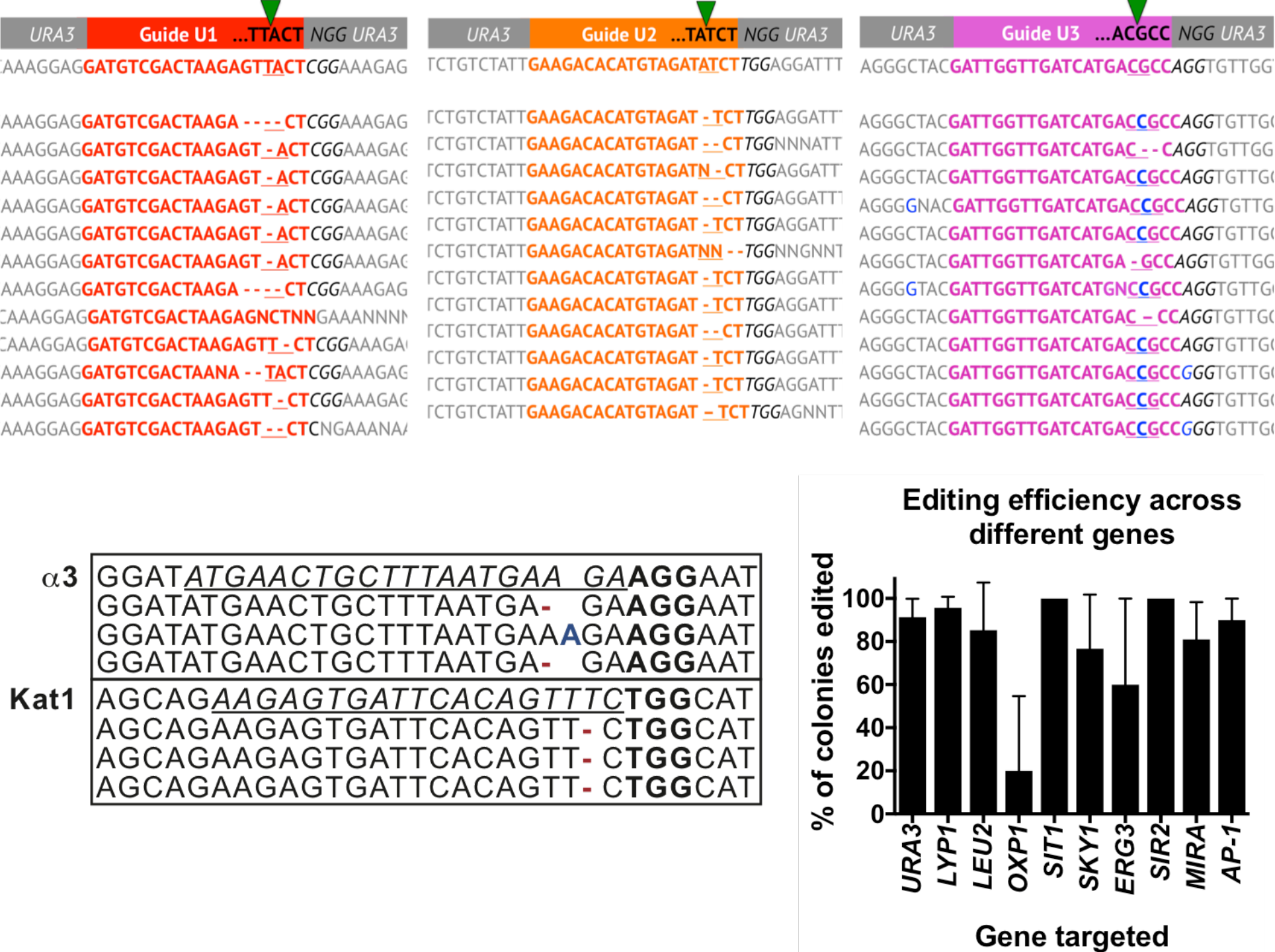
Cas9 editing outcomes and efficiency for several *K. marxianus* genes. (**A**) CRISPR_NHEJ_ results for *KmURA3* targeting experiments in the absence of donor DNA. Three regions were targeted and small insertions and deletions were found upon repair of double strand breaks by the NHEJ machinery. (**B**) Small indels can also be observed when targeting *KmKAT1* and *KmALPHA3* genes. (**C**) Editing efficiency across 10 different *K. marxianus* genes is around 75%, with some genes more editing-prone than others.

**Figure S2.**
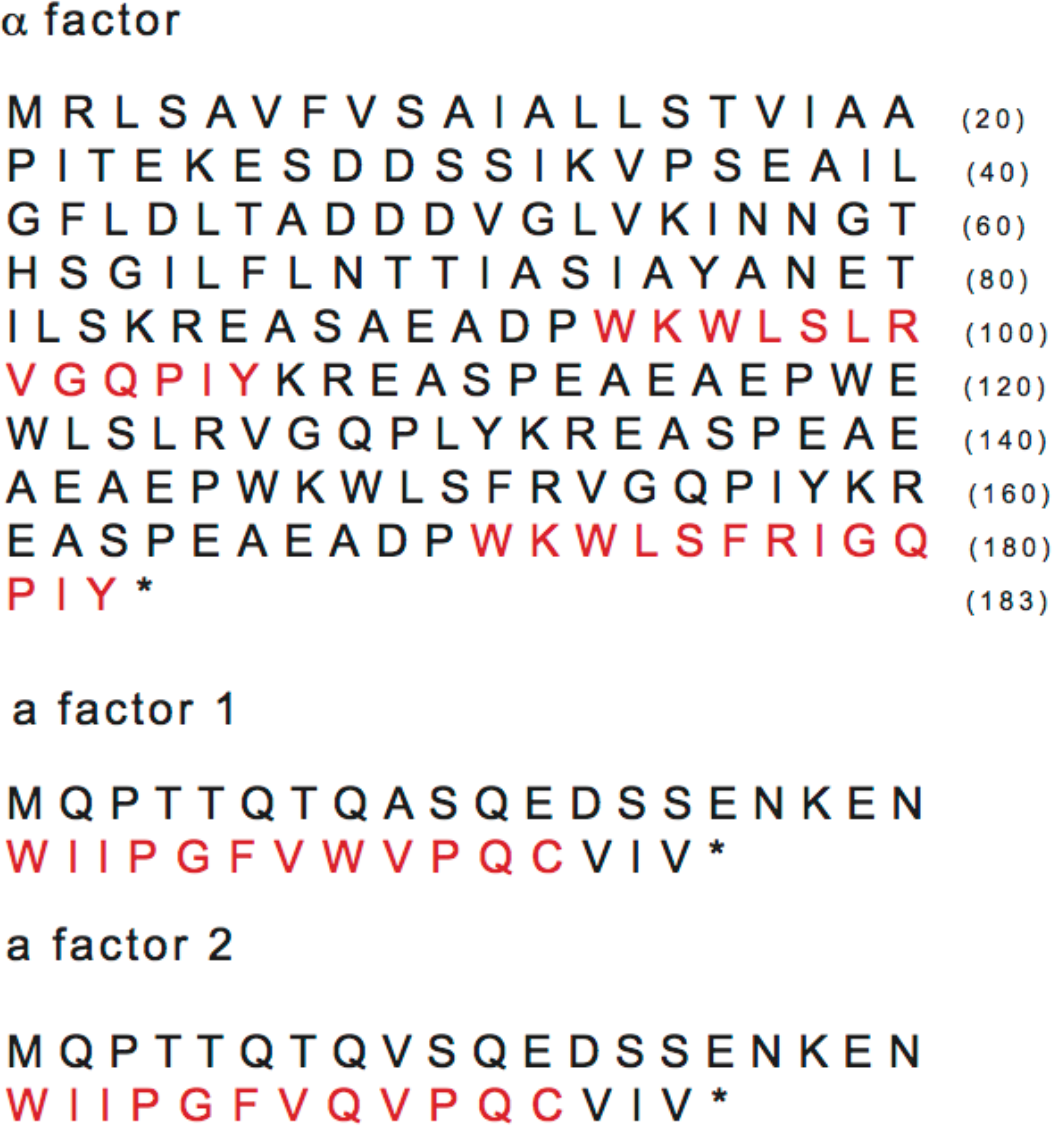
*K. marxianus* a and α factor protein sequences. Mature pheromones (in red) derive from precursor proteins, mating factor a (MFA1 and MFA2) and mating factor α (MFα1). We identified two putative *MFA* genes as well as the *MFα* gene (*KmMFαl* encoding 2 isotypes KmMFαl and KmMFα2) in the *K. marxianus* genome by reciprocal BLASTp using the *S. cerevisiae* and *K. lactis* protein sequences as queries.

**Figure S3.**
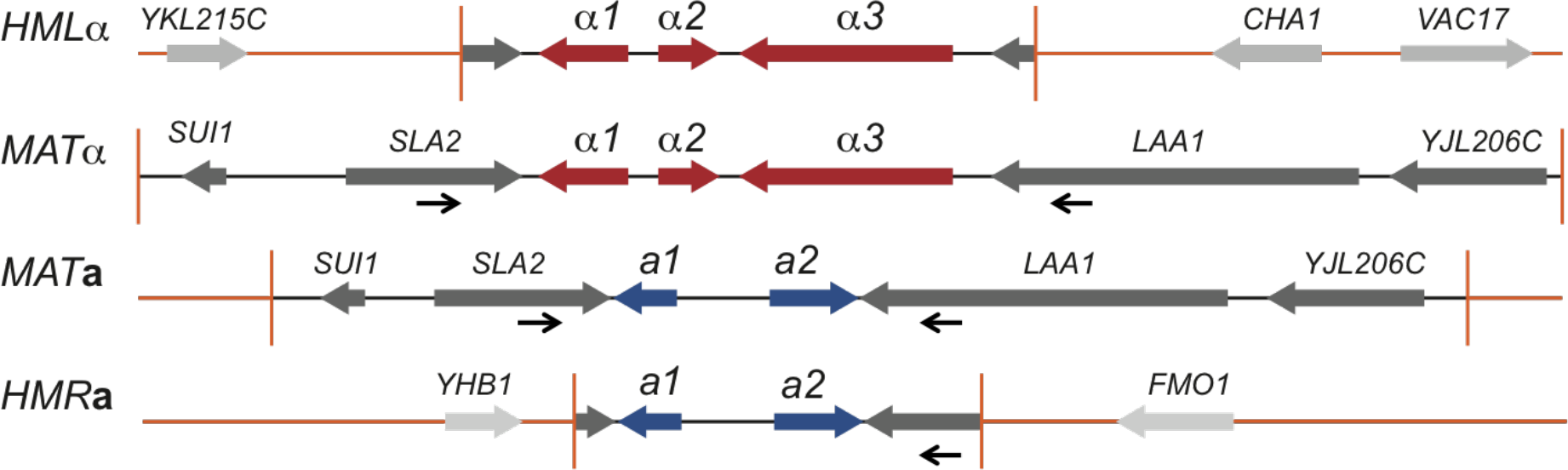
Proposed architecture of *K. marxianus MATa, MATα, HMRa* and *HMLα* loci. Black arrows indicate annealing sites for the genotyping primers (*SI Appendix*, Table S2). PCR of an **a**-type strain yield ~3480 bp product while α-type yield ~6500 bp PCR product.

**Figure S4.**
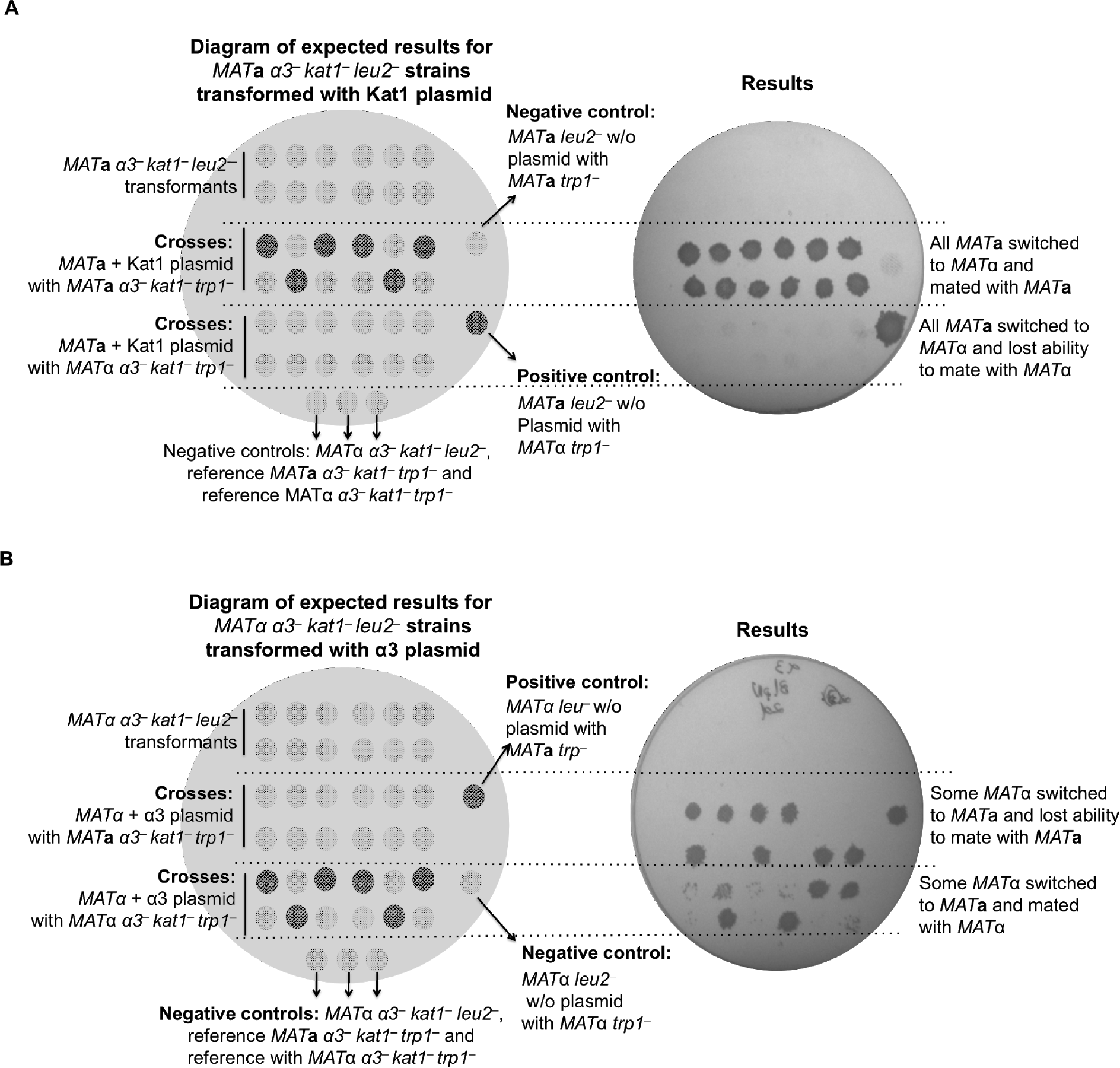
Kat1 and α*3* rescue of mating type switching. Plasmid-based ectopic expression of Kat1 and α*3* restores mating-type switching in *MAT***a** *α3*^−^ *kat1*^−^ and *MATα α3*^−^ *kat1*^−^ respectively. SCD minus (Leu, Trp) plates are shown where only diploid strains are able to grow. (**A**) Ectopic expression of Kat1 allowed a stable *MAT***a** *α3*^−^ *kat1*^−^ *leu2*^−^ strain to switch to *MAT*α and mate with the *MAT***a** *α3*^−^ *kat1*^−^ *trp1*^−^ reference strain. (**B**) Similarly, ectopic expression of α3 allowed some *MAT*α transformants to switch to *MAT***a** and mate with the *MAT*α reference strain. Note that some *MAT*α transformants did not switch to *MAT***a**, keeping the ability to mate with *MAT***a**.

**Figure S5.**
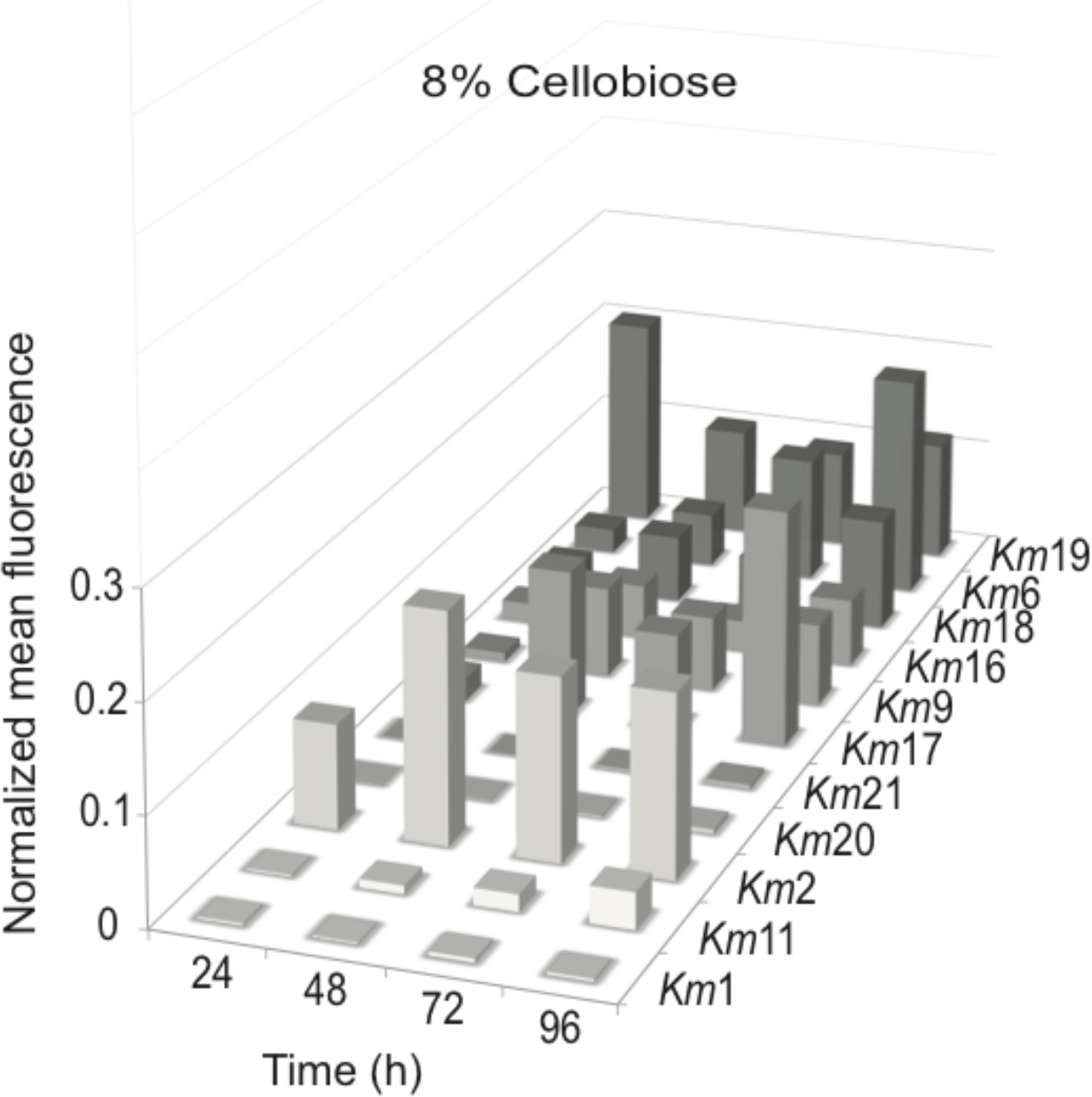
Lipogenesis of *K. marxianus* strains grown on 8% cellobiose. Normalized mean Nile red fluorescence flow cytometry of 11 wild type isolates after 24, 48, 72 and 96 hours at 42 °C in lipogenesis medium containing 8% cellobiose. Maximum values do not surpass 30% of the value obtained for the top lipid-producing strain grown on glucose.

**Figure S6.**
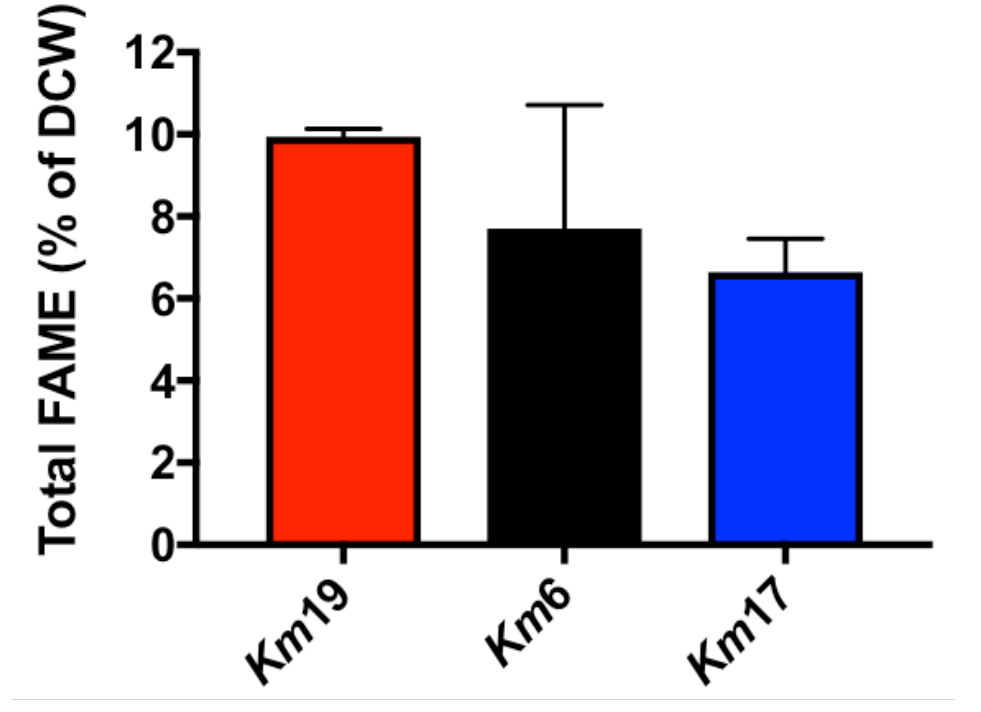
Lipid accumulation in *K. marxianus* strains. *Km*19, *Km*17 and *Km*6 percentage of fatty acids in dry cell weight after 24 hours in 8% glucose at 42 °C and 250 RPM. Lipogenesis medium contained ammonium sulfate instead of monosodium glutamate for this set of measurements. Measurements were from biological triplicates, with mean and standard deviation shown.

**Figure S7.**
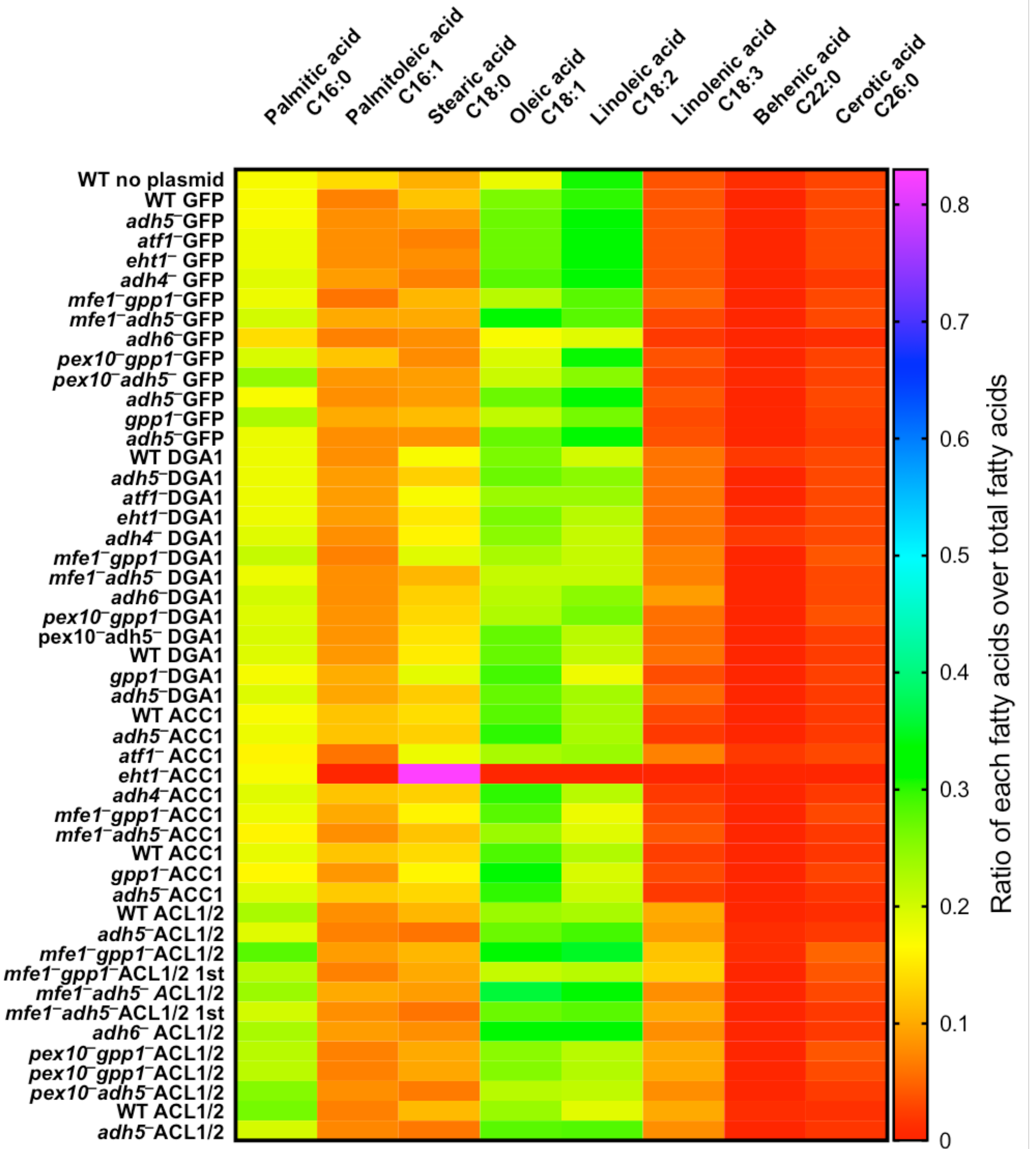
Fatty acid composition of *Km*6-derived strains. Eight fatty acids were measured using GC-FID, no appreciable difference in composition was seen among the strains except for mutant *Km*6 *eht1*^−^ bearing the ACC1 overexpression plasmid.

**Figure S8.**
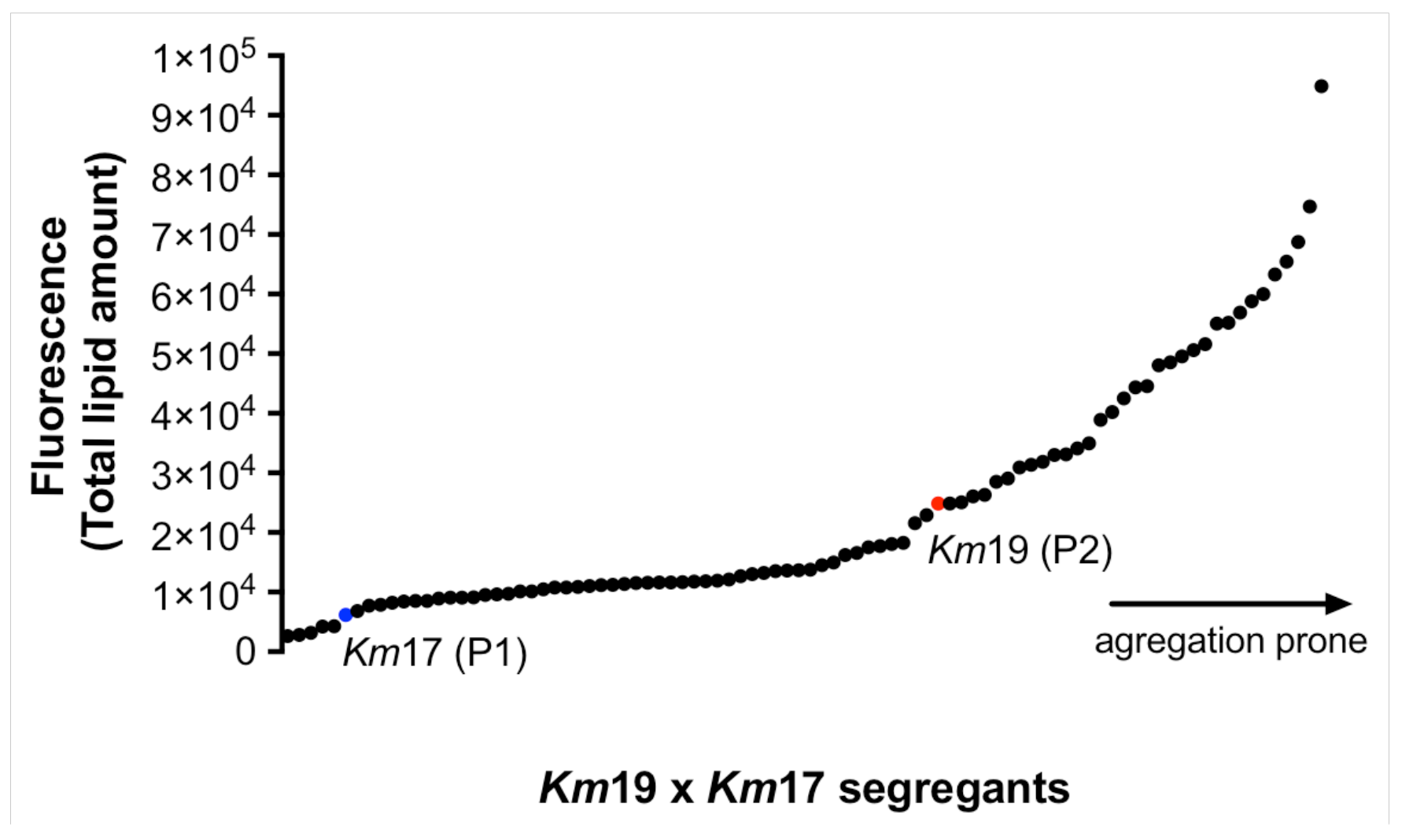
Nile red staining and flow cytometry for each single segregant from the *Km*17 × *Km*19 cross. The diploids from this cross were sporulated and spores were germinated in high temperature (44 °C). 91 spores were collected, grown in lipogenesis condition, treated with Nile red and subjected to flow cytometry.

**Figure S9.**
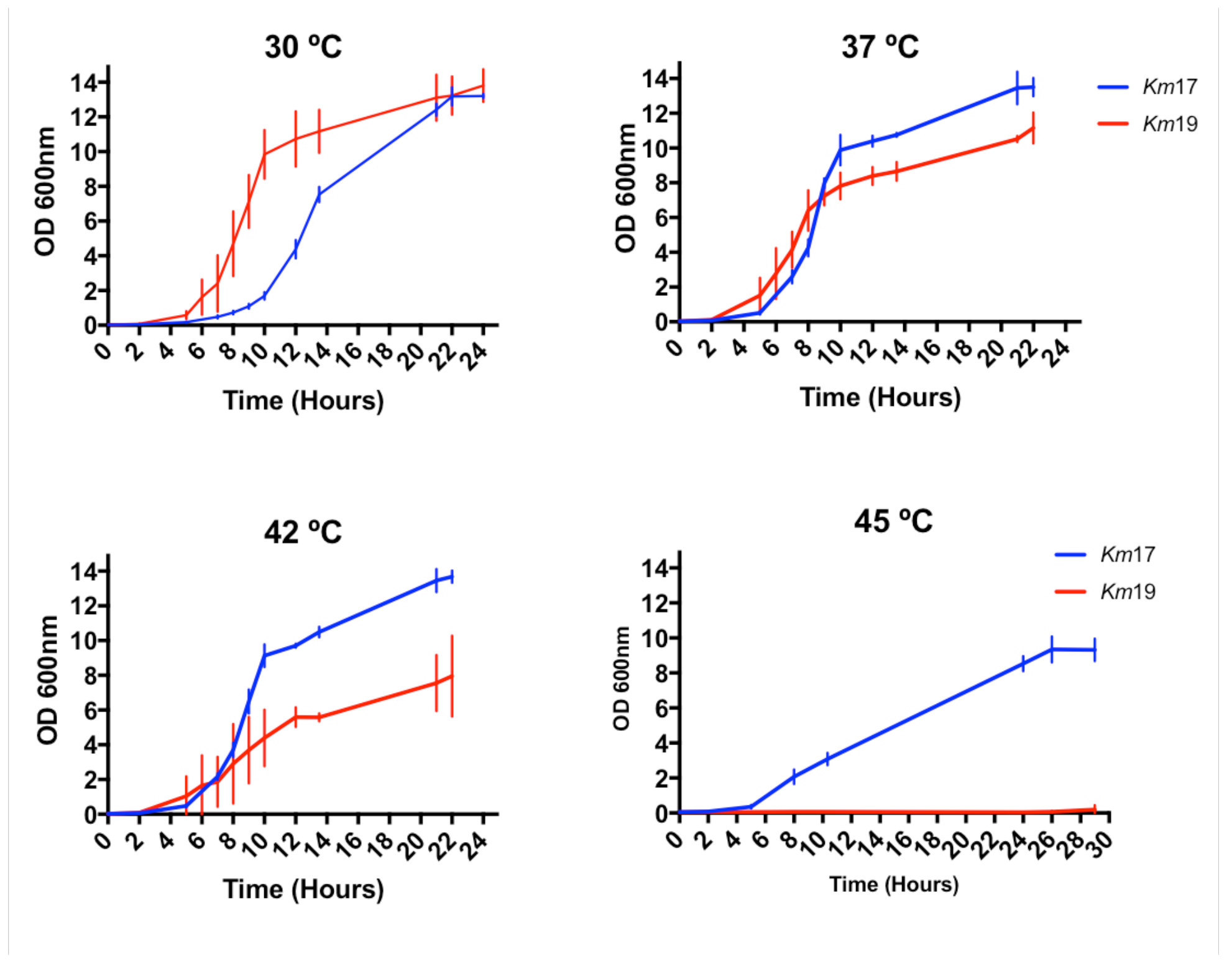
Temperature-dependence of the growth of *K. marxianus* strains tested for lipogenesis. Growth curves of *Km*17 and *Km*19 in 50 mL YPD medium at 30 °C, 37 °C, 42 °C and 45 °C and 250 RPM. Experiments are from biological triplicates.

**Figure S10.**
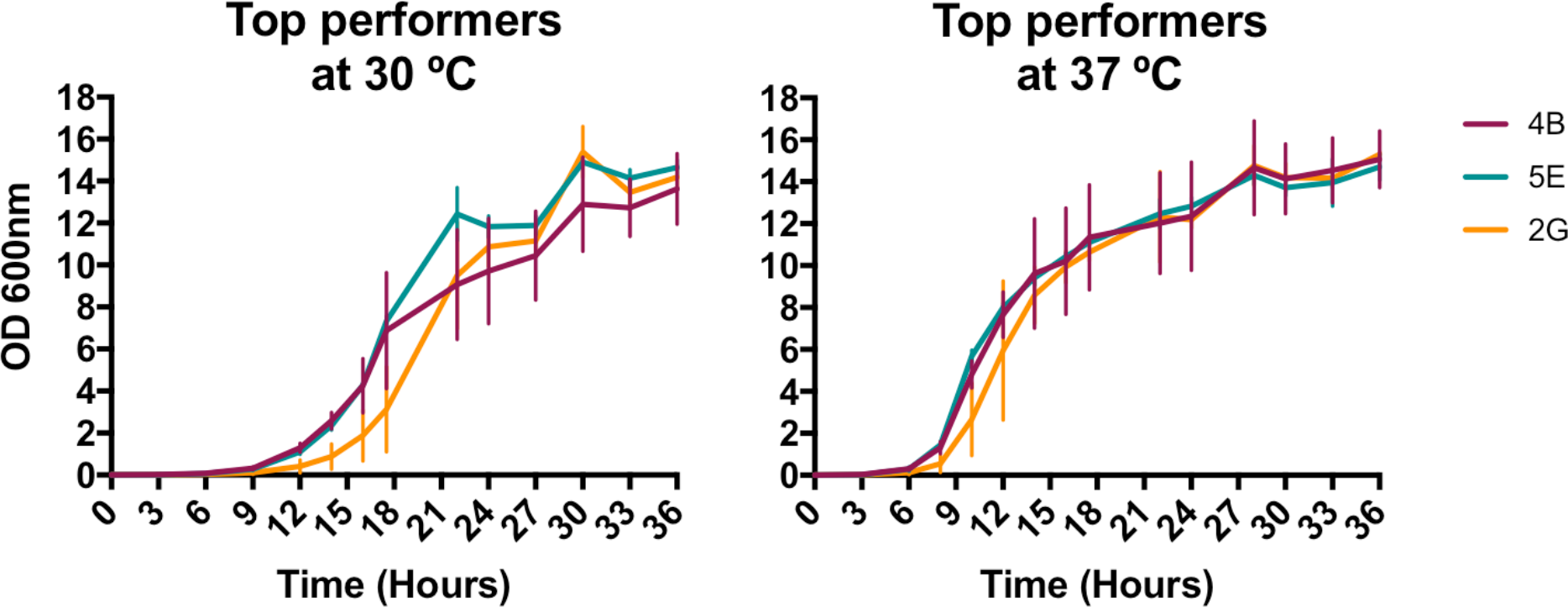
Growth curves of the highly lipogenic isolates from mating *Km*17 and *Km*19. Cells were grown in 50 mL YPD medium at 250 RPM, at 30 °C and 37 °C. Experiments are from biological triplicates.

**Table S1.**
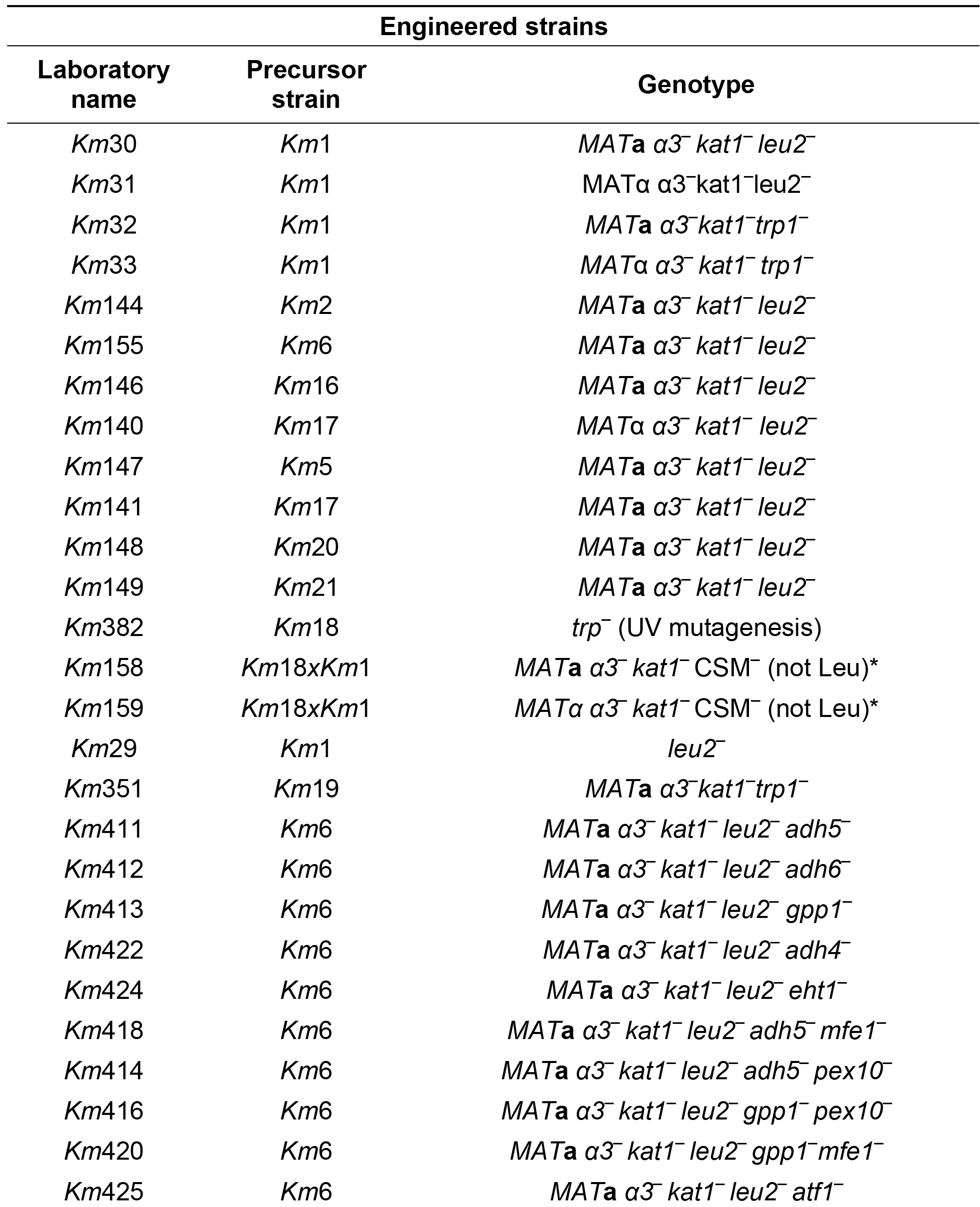
List of engineered strains.

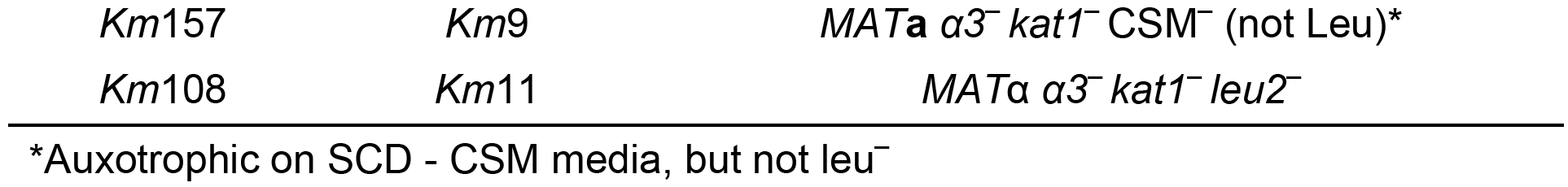

**Table S2.**
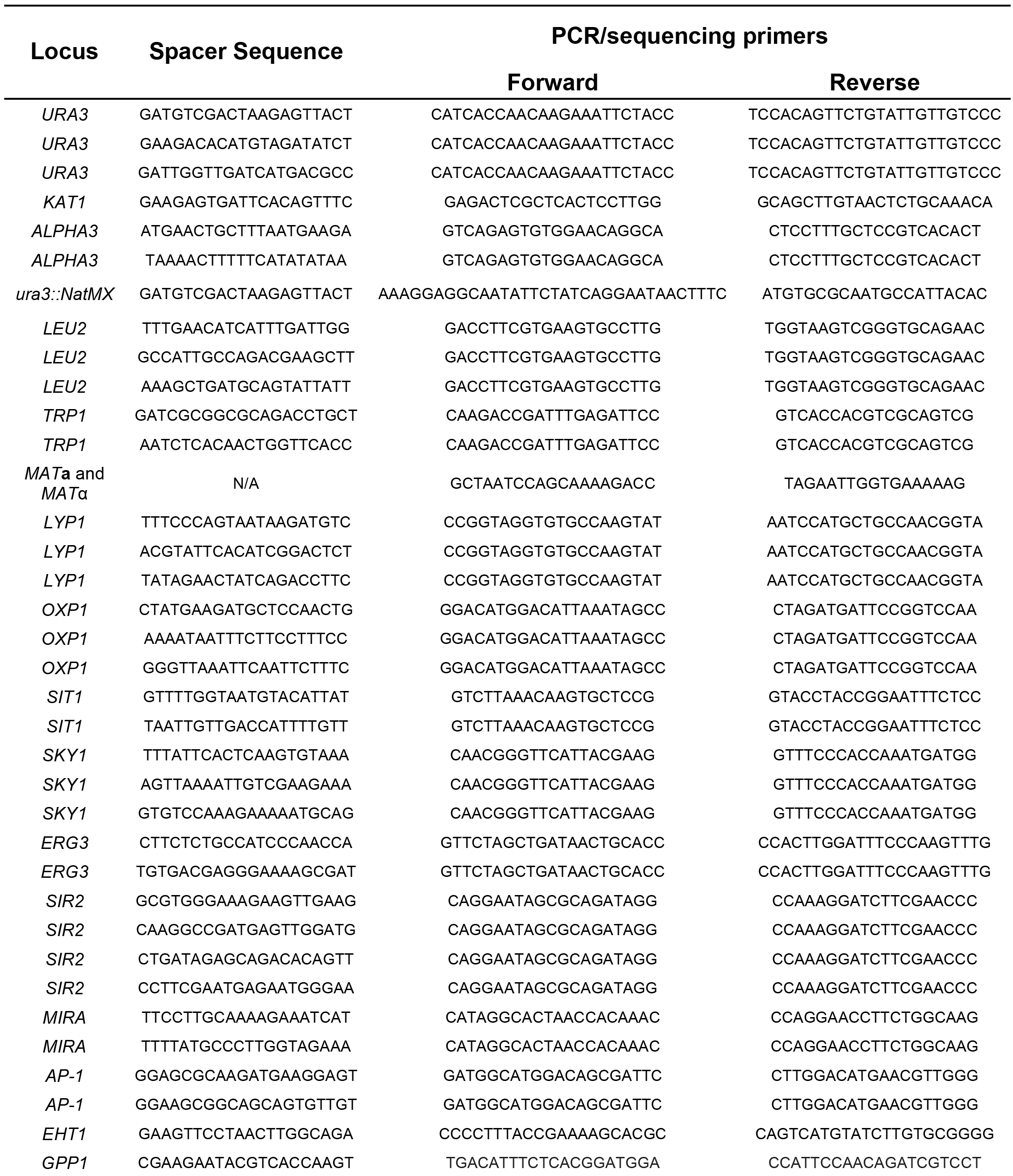
Primers and spacer sequences.

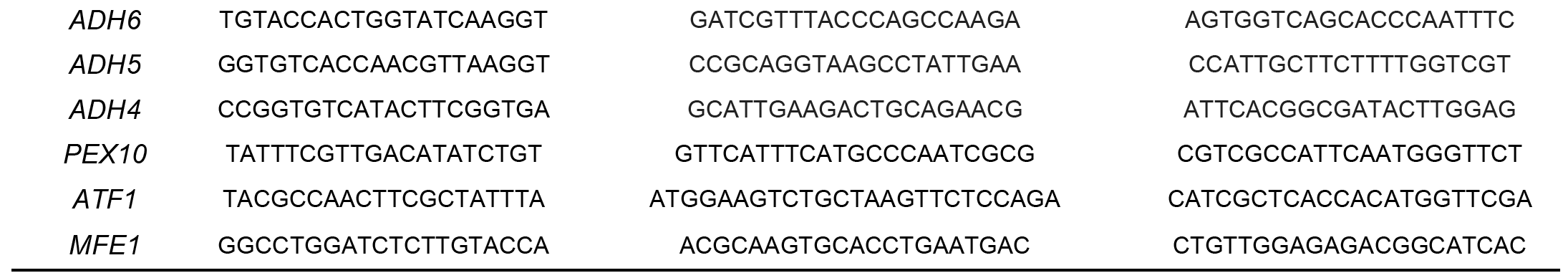

**Table S3.**
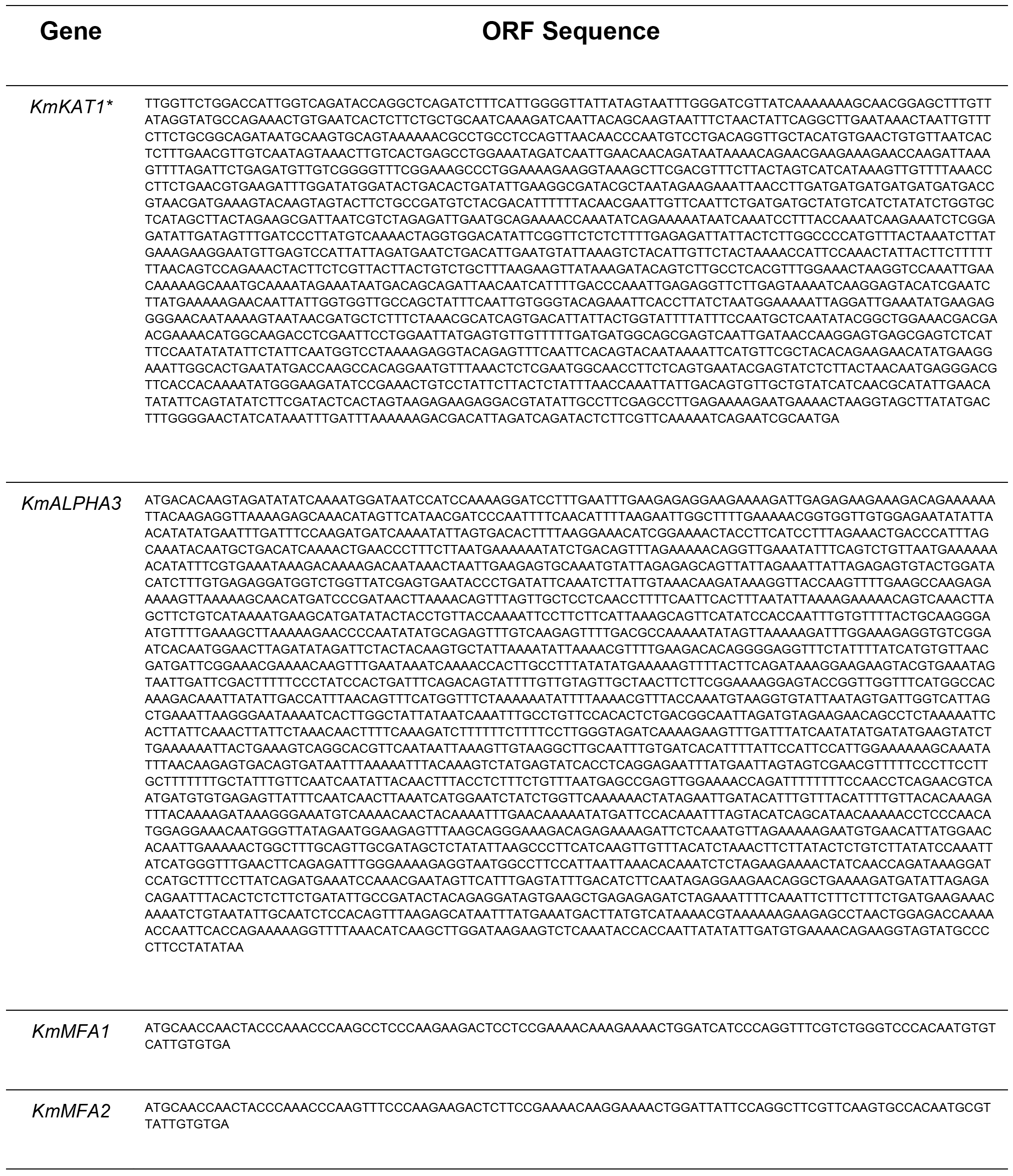
Relevant DNA sequences.

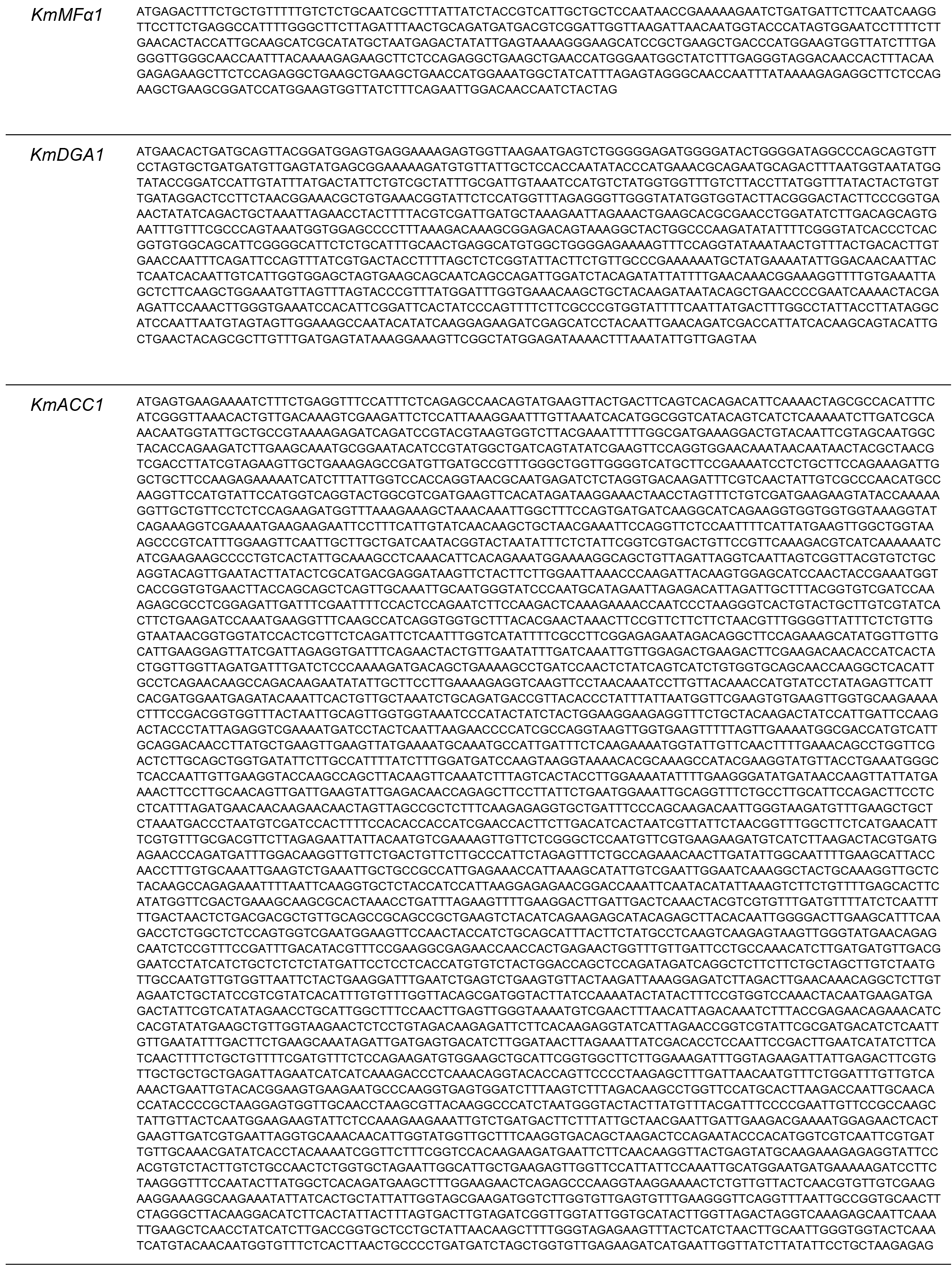

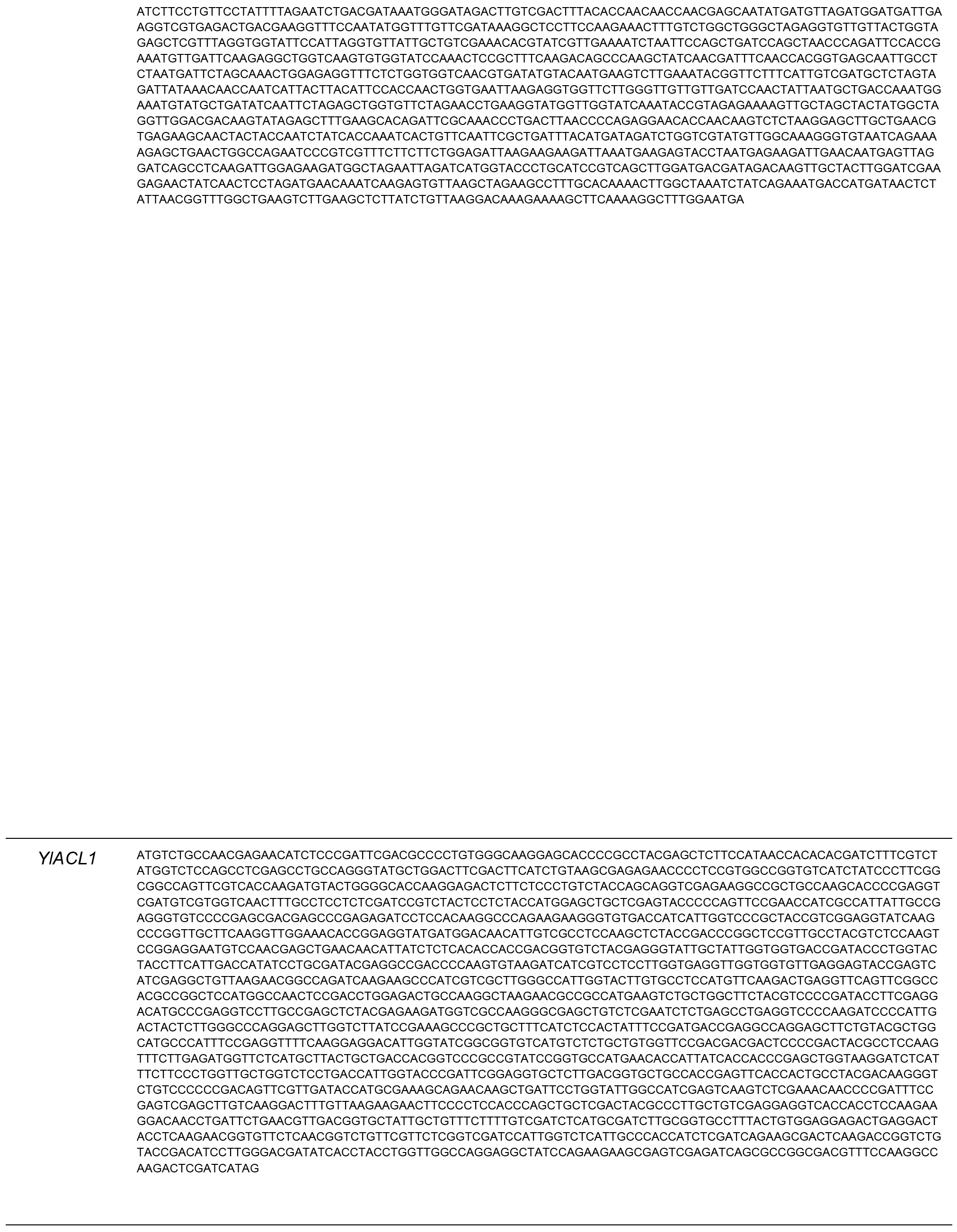

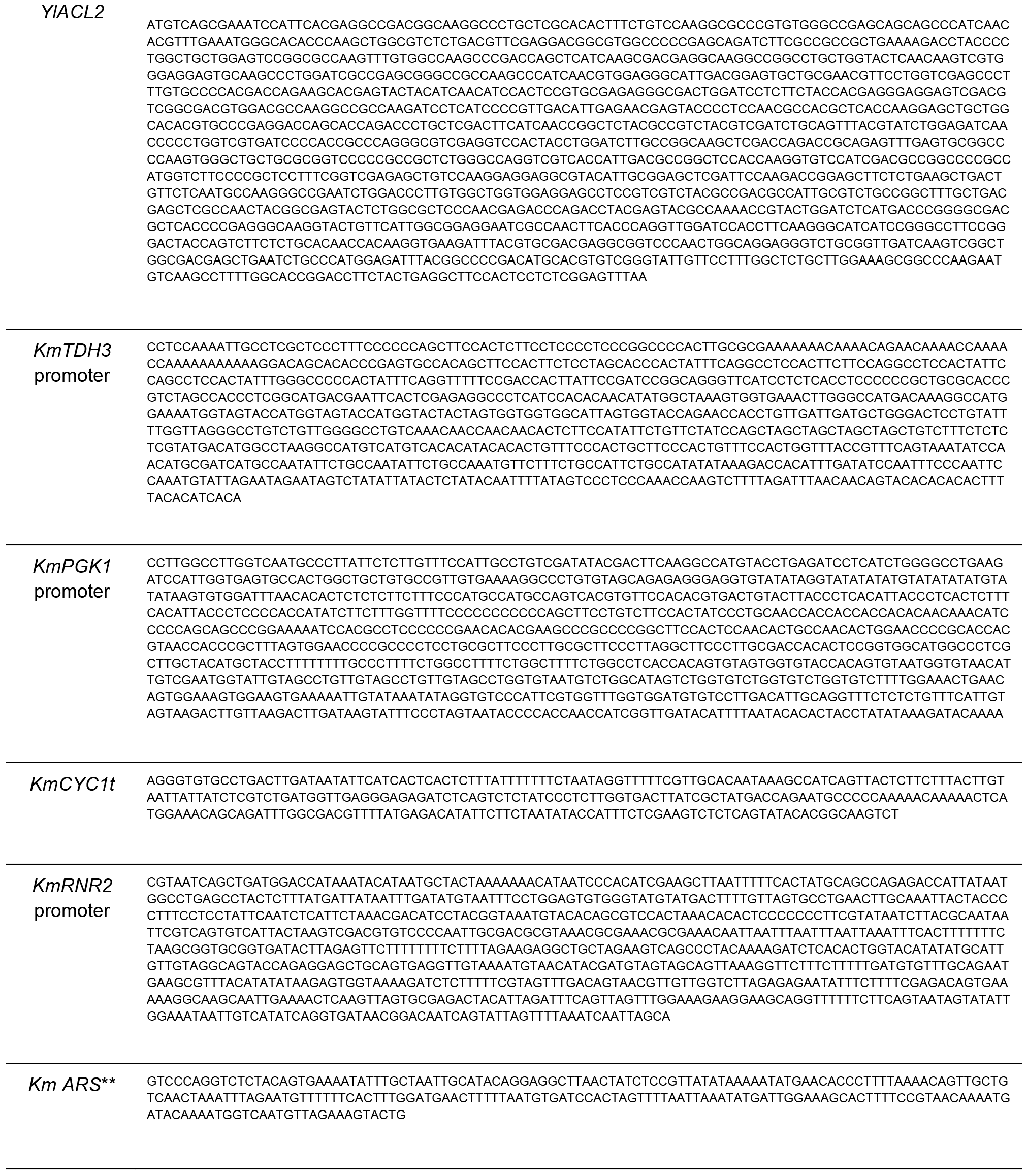

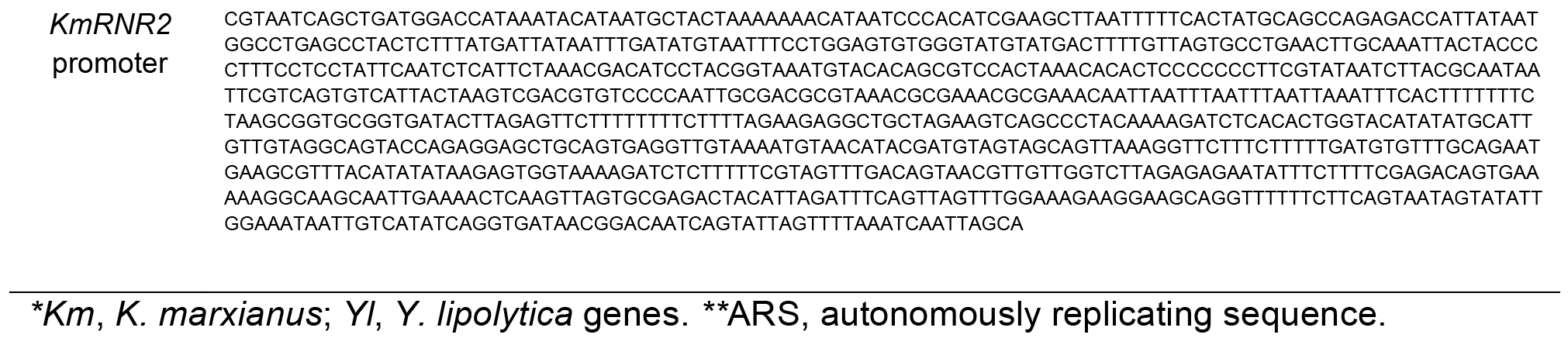

